# Immunotherapy with Pro-regenerative Macrophages from Embryonic Stem Cells Ameliorate Osteoarthritis via TNFAIP3-Mediated Chondroprotection

**DOI:** 10.64898/2025.12.12.693855

**Authors:** Zhumei Zhuang, Zicong Liu, Hang Su, Wei Sun, Qiuwen Zhu, Jingyi Xu, Jinghua Fang, Liying Li, Danni Shen, Xiaohui Zou, Hendrik Marks, Xianzhu Zhang, Hua Liu, Hongwei Ouyang

## Abstract

Immune cell-based therapies for osteoarthritis (OA) remains underexplored. Macrophages coordinate tissue repair, yet their functional heterogeneity and therapeutic applicability, particularly for designed stem cell-derived populations with tailored reparative properties are not fully delineated. Inspired by embryonic skeletal development as a paradigm of scarless regeneration, we identified a pro-regenerative macrophage population through a single-cell RNA atlas of developing skeletal tissues. We subsequently established a mechanically dynamic induction approach to efficiently generate pro-regenerative macrophages from human embryonic stem cells (hESCs), mimicking this developmental phenotype. *In vitro*, these macrophages mitigate cartilage degradation in OA explants and suppress pro-inflammatory, catabolic signaling in chondrocytes. Injection into murine OA joints robustly attenuates disease progression, reduces extracellular matrix breakdown, and resolves synovial inflammation, representing the first demonstration of immune cell-based therapy for OA. Mechanistically, these macrophages orchestrate chondroprotection not through broad immunosuppression, but by activating a TNFAIP3-dependent signaling hub that restrains inflammatory and catabolic responses in chondrocytes. This TNFAIP3 activation precisely reversed the OA-associated gene signature, suppressing catabolic mediators while enhancing reparative factors, thereby restoring the anabolic-catabolic balance. Our findings uncover a developmental blueprint for engineering pro-regenerative macrophages from stem cells and establish a pioneering targeted immune cell therapy that engages an intrinsic chondroprotective program, offering a transformative and translationally relevant strategy for OA treatment.

## Introduction

The pursuit of regenerative therapies for osteoarthritis (OA) has increasingly focused on cell-based approaches. Mesenchymal stem cell (MSC) transplantation, for instance, has demonstrated immunomodulatory effects, yet their capacity to drive robust and structural cartilage repair remains inconsistent, often failing to overcome the disease’s persistent inflammatory and catabolic milieu^1–4^. This limitation underscores the urgent need for therapeutic agents that can not only modulate inflammation but also actively and dominantly reprogram the joint environment towards a regenerative state.

Cell-based immunotherapy represents a promising frontier for OA treatment, yet its application has been hampered by a lack of defined, therapeutically potent immune cell populations. Macrophages emerge as compelling candidates given their masterful coordination of immunity, tissue homeostasis, and repair^5, 6^. However, the therapeutic potential of defined macrophage populations remains largely unecharted, primarily due their profound functional heterogeneity. This heterogeneity manifests in two key dimensions: developmental origin and functional polarization. In terms of origin, tissue macrophages consist of embryonically derived tissue-resident macrophages (TRMs) and those arising from recruited monocytes (monocyte-derived macrophages, MDMs)^7^. TRMs are critical for developmental morphogenesis and maintaining tissue homeostasis, often exhibiting stable, pro-reparative phenotypes^8^. MDMs infiltrate upon injury and can adopt context-dependent, often inflammatory or fibrogenic roles ^9, 10^. Beyond origin, macrophage functional states extend far beyond the classical M1/M2 spectrum, comprising distinct subsets with diverse functional fates. Although specific macrophage phenotypes, particularly pro-resolutive ones, have been linked to repair outcomes^11–14^. the field still lacks a consensus definition of a universally pro-regenerative human macrophage and a scalable method to generate it. This critical gap severely hinders the development of reliable macrophage-based immunotherapies.

We posited that the blueprint for such a regenerative macrophage resides in embryogenesis, a period of flawless tissue development and scarless healing. By computationally deconstructing a published single-cell atlas of developing skeletal tissue^15^, we revealed that embryonic macrophages are strategically positioned and engage in active crosstalk with skeletal stem cells and chondroprogenitors, suggesting a master regulatory function in coordinating chondrogenesis. This discovery established a developmentally inspired rationale for harnessing specific embryonic macrophage phenotypes. Inspired by this blueprint, we sought to recapitulate this macrophage regenerative identity *in vitro* using human pluripotent stem cells (hPSCs). This category encompasses both human embryonic stem cells (hESCs) and human induced pluripotent stem cells (hiPSCs). These cells possess the dual attributes of self-renewal and multipotency, which enable their directed differentiation to yield a theoretically unlimited source of macrophages^16, 17^. While several methods for differentiating PSCs into macrophages (PSC-Macs) *in vitro* have been described^18–21^. Macrophages derived via classical protocols are often time-consuming, inefficient and ill-defined; they frequently constitute heterogeneous populations that resemble fetal or inflammatory states, lacking a stable functional identity. This uncertainty over whether PSC-Macs represent a distinct, therapeutically relevant subtype remains a major barrier to their clinical translation.

To address this, we established a novel, stage-adapted differentiation strategy to produce human embryonic stem cell-derived macrophages (hESC-Macs) exhibiting a stable pro-regenerative phenotype. Comparative transcriptomic profiling revealed that our hESC-Mac conserved a pro-regenerative gene signature homologous to their embryonic skeletal counterparts. A significant fraction of hESC-Macs co-expressed a suite of genes associated with tissue repair, among which the Mannose Receptor 1 (MRC1/CD206) and High Mobility Group Box 1 (HMGB1) genes were among the most robustly and consistently elevated markers within this signature.

Here, we demonstrate that these developmentally inspired hESC-Mac exert potent chondroprotective and reparative effects in both human OA chondrocyte cultures and a murine OA model. Mechanistically, this occurs primarily through the precise activation of a TNFAIP3-dependent chondroprotective program in chondrocytes, which serves as a master regulator to suppress the destructive NF-κB axis. Our work identifies a targetable regenerative program for generating potent, macrophage-based immunotherapies for osteoarthritis.

## Results

### Decoding embryonic macrophage heterogeneity reveals functional specialization and chondro-regenerative potential

To characterize macrophage populations during human skeletal development, we integrated and re-analyzed published single-nucleus RNA sequencing (snRNA-seq) data comprising 372,937 cells from knee and shoulder joint specimens obtained from twelve donors ranging from 5 to 18 post-conception weeks (PCW)^15^. Unbiased clustering identified eight major cellular compartments, with mesenchymal and immune cells exhibiting distinct distribution patterns across developmental stages and anatomical regions **(Supplementary Fig. 1a, b)**. Further subclustering of the immune compartment revealed that macrophages constituted a dominant population with previously unappreciated heterogeneity **(Supplementary Fig. 1c, d)**. We classified embryonic skeletal macrophages into five transcriptionally distinct clusters: LYVE1□, COL1A1□, COL9A1□, PAX7□, and CD34□. LYVE1□ macrophage was the most abundant, comprising approximately 60% of the population across anatomical sites. COL1A1□ and COL9A1□ macrophages were present at moderate frequencies, while PAX7□ and CD34□ subsets were less frequent **(Supplementary Fig. 1e, f)**.

Transcriptomic profiling revealed subset-specific gene signatures suggestive of functional specialization. LYVE1□ macrophages expressed genes associated with immunomodulation and tissue homeostasis, including TIMD4, LYVE1, MRC1, CD163, VEGFA and VSIG4, consistent with an M2-like polarization state. COL1A1□ macrophage displayed elevated expression of fibrillar collagens (COL3A1, COL5A1), the matricellular protein SPARC, and secreted factors including SPP1, IGFBP5, and IGFBP7, suggesting roles in fibrotic matrix organization. In contrast, COL9A1□ macrophages expressed genes characteristic of hyaline cartilage, including COL9A1, COL9A2, COL2A1, ACAN, and the chondrogenic transcription factor SOX9. CD34□ macrophages expressed a gene set associated with hematopoietic niche function (CD34, VCAM1, CXCL12, TEK), while PAX7□ macrophages were enriched for genes involved in proliferation and metabolism (CECR2, SLC38A1), suggesting a progenitor-like state **(Supplementary Fig. 1g)**. Pseudotemporal analysis revealed a structured developmental hierarchy. LYVE1□ macrophages appeared earliest, present from 5 PCW and persisting through 11 PCW. COL1A1□ and COL9A1□ macrophages emerged subsequently, followed by PAX7□ and CD34□ clusters around 8 PCW **(Supplementary Fig. 1h, i)**. This timing indicates that macrophages colonize the embryonic skeletal niche and acquire subset-specific identities prior to the initiation of robust chondrogenesis at approximately 8 PCW **(Supplementary Fig. 1j)**.

To assess potential macrophage-chondrocyte communication, we performed ligand-receptor interaction analysis focusing on pathways implicated in chondrogenesis and cartilage homeostasis. This analysis identified extensive predicted interactions between macrophage subsets and mesenchymal lineages, including chondroprogenitors and chondrocytes (**Supplementary Fig. 2 and 3a**). Macrophages were predicted to serve as sources of TGF-β, PDGF, BMP, and IL-10 signaling, with subset-specific patterns. LYVE1□ macrophages showed high expression of IL10 and predicted interactions with chondrocytes via TGF-β pathway components (**Supplementary Fig. 3b, c**). COL9A1□ macrophages expressed ligands for VEGF and TGF-β pathways, while CD34□ macrophages showed predicted CXCL-mediated interactions (**Supplementary Fig. 3d**). While these computational predictions require functional validation, they suggest that embryonic macrophage subsets may contribute to the signaling environment during skeletal development.

### A biomimetic dynamic platform efficiently generates hESC-derived macrophages with high yield and purity for therapeutic application

Based on our observation that embryonic macrophages with potential regenerative properties are present during skeletal development, we sought to develop a robust system for generating similar macrophages from human embryonic stem cells (hESCs) *in vitro*. We established a directed, stepwise differentiation protocol that recapitulates key stages of primitive hematopoiesis and myelopoiesis, creating what we term a “biomimetic developmental paradigm” for macrophage production. Building upon a conventional static suspension method^22^, we developed a dynamic shaker-based culture system to enhance hematopoietic induction and myeloid lineage commitment. This approach achieved myeloid lineage specification within 6 days, compared to 10 days for the static method—a 40% reduction in timeline **(Fig. 1a)**. The differentiation protocol comprises two phases: an initial suspension culture period followed by adherent culture, with hESC-derived macrophages (hESC-Mac) emerging as early as day 10 **(Fig. 1b)**. Cells displayed characteristic macrophage morphology and expressed the pan-macrophage marker CD68 by immunofluorescence **(Fig. 1c)**. Flow cytometric analysis demonstrated progressive upregulation of CD45 and CD14, with over 90% of cells co-expressing these markers by day 20, indicating efficient macrophage differentiation **(Fig. 1d)**. Quantitative RT-PCR confirmed stage-appropriate transcriptional dynamics: silencing of pluripotency genes (POU5F1, SOX2), transient activation of mesodermal markers (MIXL1, FLK1), sequential expression of hematopoietic (RUNX1, CD34) and myeloid (SPI1, CEBPA) genes, and sustained expression of macrophage markers (CSF1R, CX3CR1) **(Fig. 1e)**.

**Figure 1.**
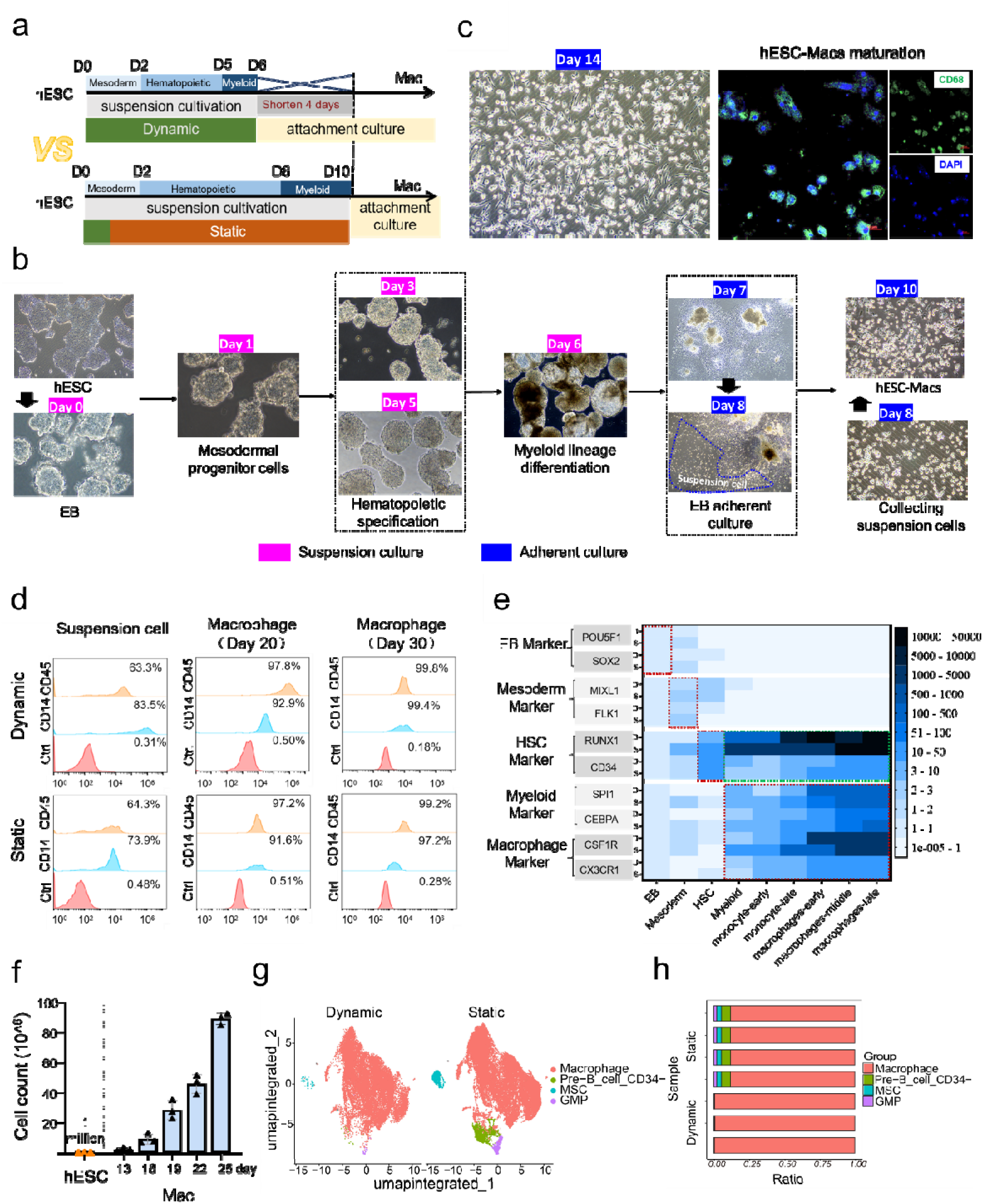
Efficient generation and characterization of hESC-Mac through a mechanically dynamic differentiation system. (a) Schematic comparison of conventional (lower) and optimized (upper) differentiation protocols. The conventional static suspension culture method requires 10 days to generate myeloid progenitor stages. The optimized dynamic culture system shortens the differentiation process by 4 days through improved hematopoietic specification and myeloid commitment. (b) Bright-field microscopy images showing the morphological progression of cells at critical time points during dynamic differentiation, from hESC stage through mesoderm, hematopoietic, and macrophage commitment. Suspension culture (red), Adherent culture (blue), Scale bars=100 μm. (c) Immunofluorescence analysis confirming the successful generation of hESC-Mac on Day 14 of differentiation. CD68 immunofluorescence staining (red) confirms macrophage identity. Nuclei are visualized with DAPI (blue). Scale bars, 50 μm. (d) Flow cytometry analysis of collecting suspension cells, cultured macrophages at day 20 and day 30, demonstrating co-expression of the macrophage surface antigens CD14 and CD45. The percentage represents the purity of macrophage. (e) RT-qPCR analysis of the key developmental stages and corresponding marker genes during the differentiation process: embryoid body (POU5F1, SOX2), mesoderm (MIXL1, FLK1), hematopoietic (RUNX1, CD34), myeloid (SPI1, CEBPA), and macrophage (CSF1R, CX3CRI) lineages. (f) Quantification of hESC-Mac yield over time in the dynamic culture system, revealing a progressive increase from Day-13 to approximately 10□ cells per differentiation by Day-25 (n=3 independent differentiations; each dot = one sample; data are mean ± SD). scRNA-seq validation of hESC-Mac identity and purity. (g) UMAP projection of cells from our dynamic differentiation culture (hESC-Mac) and a control static culture, colored by annotated cell types. (h) Proportional composition of different cell types identified in hESC-Mac versus the static control culture, bar indicates replicates, color indicates cell types.

hESC-Mac displayed low surface expression of MHC class I (HLA-A, B, C) and MHC class II (HLA-DR, DP, DQ) molecules **(Supplementary Fig. 4a)**, a favorable trait for potential therapeutic applications. To rigorously validate the embryonic character of our hESC-Mac, we conducted single-cell RNA sequencing and merged our dataset with publicly available profiles of yolk sac^23^and and placenta-derived macrophages^24^. Unsupervised clustering and UMAP visualization revealed that hESC-Mac shared transcriptional signatures with primary embryonic macrophage subsets, confirming their authentic developmental origin and functional relevance **(Supplementary Fig. 4b-d)**.

Beyond developmental fidelity, our system addressed critical translational limitations. The dynamic differentiation platform also achieved substantial scalability, with yields increasing steadily to approximately 100 million macrophages per differentiation by day 25 **(Fig. 1f)**, addressing a critical limitation of donor-dependent primary macrophages. Single-cell transcriptomic analysis confirmed the homogeneity of the final product, with nearly 100% of cells clustering as macrophages, a significant improvement over static control cultures, which contained contaminating populations such as pre-B cells, mesenchymal stromal cells, and granulocyte-monocyte progenitors (**Fig. 1g, h**). We hypothesize that the dynamic differentiation strategy of hESCs may promote the expression of macrophage-related genes by activating mechanosensitive signaling pathways (evidenced by the high expression of CTNNB1 (β-catenin)), remodeling the cytoskeleton dynamics (involving ACTB, RHOA, and ROCK2), and facilitating the nuclear translocation of the transcription factor IRF8 **(Supplementary Fig. 4e)**. However, the precise underlying mechanism requires further investigation.

In summary, we have established a biomimetic platform that not only accelerates macrophage generation but also faithfully recapitulates embryonic skeletal development. The combination of high yield, exceptional purity, low immunogenicity, and embryonic-like characteristics positions these hESC-Mac as an ideal cell source for regenerative applications, particularly for cartilage repair where embryonic macrophages demonstrate inherent therapeutic potential.

### hESC-Mac exhibit a unique pro-regenerative phenotype orchestrated by MRC1 and HMGB1 co-expression

Following the establishment of a robust and efficient platform for hESC-Mac generation, we next aimed to comprehensively define their molecular identity and functional potential at the transcriptional level. To this end, we first subjected hESC-Mac to single-cell RNA sequencing (scRNA-seq) analysis, which enabled the unbiased partitioning of these cells into 9 distinct subpopulations (subclusters 0–8) based on transcriptional heterogeneity (**Fig. 2a**). To gain insights into the *in vivo* relevance of hESC-Mac, we performed pseudo-bulk Spearman correlation analysis to compare the transcriptional profiles of hESC-Mac subpopulations with those of previously identified macrophage subsets isolated from embryonic skeletal tissue. We discovered that hESC-Mac correlated most strongly with LYVE1^+^ macrophages, a dominant subset constituting ∼60% of embryonic skeletal macrophages, suggesting a shared core transcriptional program (**Supplementary Fig. 5a**).

**Figure 2.**
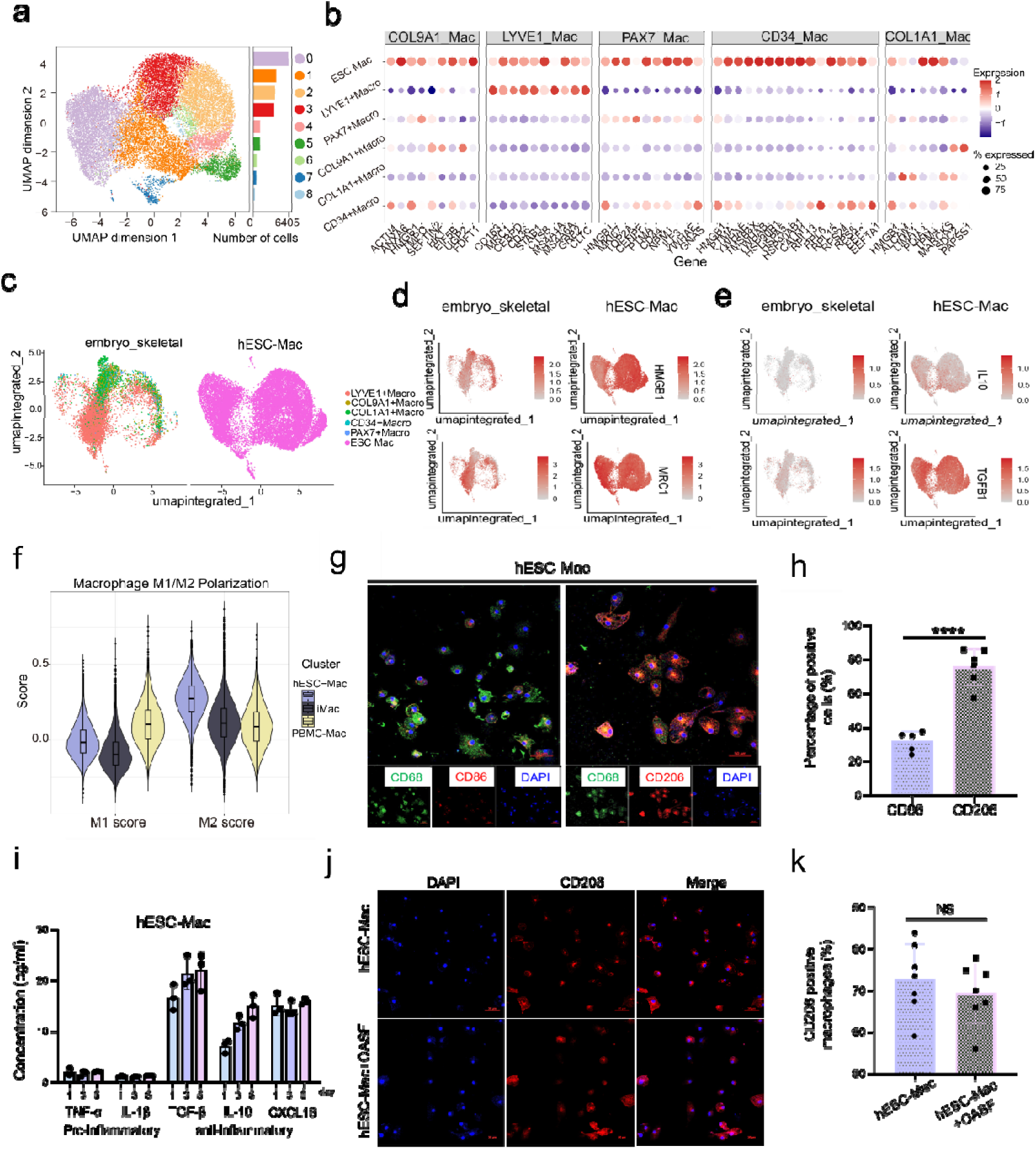
hESC-Mac recapitulates a pro-regenerative embryonic macrophage phenotype characterized by MRC1 and HMGB1 co-expression. (a) UMAP visualization of hESC-Mac. Points represent cells, with color indicating local cell density across the two primary UMAP dimensions, showing nine distinct subpopulations. (b) Dot plot depicting the co-expression patterns of key functional genes between hESC-Mac and specific embryonic macrophage subsets (LYVE1^+^ Macro, COL9A1^+^ Macro, PAX7^+^ Macro, CD34^+^ Macro, COL1A1^+^ Macro). (c) UMAP projection integrating single-cell RNA-seq profiles of primary embryonic skeletal macrophages with hESC-Macs, showing their close transcriptional relationship. (d) UMAP plot shows the expression of representative pro-regenerative and anti-inflammatory markers (MRC1, HMGB1) and (e) (IL10, TGFB1) in the integrated dataset, color indicates log-normalized expression value. (f) M1 and M2 polarization scores analysis for hESC-Mac, iMac (static protocol), and PBMC-Mac, based on M1 and M2 markers, demonstrating a uniquely skewed M2-high/M1-low phenotype in hESC-Mac. (g) Representative immunofluorescence images and (h) corresponding positive quantification of the anti-inflammatory marker CD206 (MRC1) and the pro-inflammatory marker CD86 in CD68-labeled hESC-Mac. Scale bar, 50 μm. (i) ELISA quantification of pro-inflammatory cytokines (TNF-α and IL-1β) and anti-inflammatory cytokine (TGF-β, IL10 and CXCL18) secretion in hESC-Mac. Representative immunofluorescence images (j) and quantification (k) of CD206 (MRC1) expression in hESC-Mac following treatment with osteoarthritic synovial fluid (OASF) or control medium, demonstrating phenotype stability under pathological challenge. Scale bar, 50 μm. Data are represented as mean ± SEM. *P<0.05, **P<0.01, ***P<0.001, ****P<0.0001; NS: No significant.

Analysis of shared gene expression patterns revealed that hESC-Mac and embryonic macrophage subsets co-expressed genes associated with distinct functional programs. hESC-Mac shared expression of pro-repair (CD163, MRC1 and CEBPD) and efferocytosis/phagocytosis (CD36, STAB1 and CTSB) genes with LYVE1^+^ macrophages. These cells simultaneously expressed transcripts encoding the damage-sensing and repair-initiating factors HMGB1 and ACTN4, as well as matrix synthesis and assembly (EIF5B, UGP2 and FDFT1) with COL9A1^+^ macrophages. A pro-proliferative signature (MKI67, TOP2A and CENPF), indicative of a microenvironment conducive to tissue expansion, was shared with PAX7^+^ macrophages. Co-expression with CD34^+^ macrophages included genes for initiating repair (HMGB1 and TMSB4X) and providing a supportive niche (PTMA, YWHAB and YWHAE), alongside cytoskeletal stabilization genes (LIMA1 and TPM4) (**Fig. 2b**). This comprehensive co-expression profile indicates that hESC-Mac possesses a multifaceted phenotype consistent with specialized tissue-repair macrophages. Single-cell RNA sequencing integration with embryonic macrophages confirmed these findings at cellular resolution, demonstrating close transcriptional relationships (**Fig. 2c**). hESC-Mac specifically upregulated key markers of anti-inflammation (MRC1 and CD163), damage clearance and repair initiation (CD36 and HMGB1), tissue remodeling (ACTN4 and ALCAM), and progenitor pool expansion (MKI67, CD44 and PTMA) (**Fig. 2d and Supplementary Fig. 5b**). Notably, elevated expression of upstream regulators IL10 and TGFB1 revealed reinforced anti-inflammatory and pro-reparative transcriptional circuits (**Fig. 2e**).

To benchmark these properties against conventional sources, we then compared hESC-Mac with macrophages from the standard static protocol (iMac) and from peripheral blood mononuclear cells (PBMC-Mac). Integrated scRNA-seq analysis confirmed the superior purity of hESC-Mac cultures (**Supplementary Fig. 6a, b**) and revealed their unique enrichment for anti-inflammatory (MRC1, IL10 and STAT6) and pro-regenerative (TGFB1 and PDGFB) factors (**Supplementary Fig. 6c**). Polarization assessment definitively established that hESC-Mac exhibit the highest M2 and lowest M1 scores, defining a uniquely skewed pro-repair phenotype distinct from both control groups (**Fig. 2f and Supplementary Fig. 6d**).

We functionally validated this anti-inflammatory predisposition through multiple approaches. Immunofluorescence staining revealed that ∼70% of hESC-Mac expressed CD206 (encoded by MRC1), a frequency significantly higher than PBMC-Mac (**Fig. 2g, h and Supplementary Fig. 7a, b**). Consistently, ELISA detected significantly elevated secretion of anti-inflammatory and pro-repair mediators (TGF-β1, IL-10 and CXCL18) from hESC-Mac (**Fig. 2i**), unlike the PBMC-Mac (**Supplementary Fig. 7c**). This reparative phenotype remained stable under pathological challenge, as treatment with osteoarthritic synovial fluid (OASF) did not significantly alter the high anti-inflammatory marker expression in hESC-Mac (**Fig. 2j, k**). Furthermore, these anti-inflammatory hESC-Mac retained robust phagocytic capacity (**Supplementary Fig. 7d**), indicating their functional competence is not compromised by their reparative skew.

Overall, based on the above data, it can be demonstrated that our biomimetic platform successfully generates hESC-Mac with a stable, pro-regenerative phenotype defined by MRC1 and HMGB1 co-expression. This establishes a reliable cellular product for investigating and deploying macrophage-based immunotherapy.

### hESC-Mac alleviates cartilage degeneration *in vitro*

Equipped with a defined population of pro-regenerative hESC-Mac, we next evaluated their therapeutic potential in human OA systems, testing the central premise of our study: whether developmentally inspired macrophages can effectively treat osteoarthritis. Using patient-derived tissues and multiple experimental paradigms, we systematically investigated their chondroprotective potential, with PBMC-derived macrophages (PBMC-Mac) serving as reference controls throughout. Initial evaluation using cartilage explants from OA patients revealed that co-culture with hESC-Mac for 7 days significantly enhanced glycosaminoglycan (GAG) content compared to both untreated controls and PBMC-Mac co-cultures, indicating superior extracellular matrix preservation—a critical indicator of cartilage health (**Fig. 3a, b**). To determine whether these protective effects extended to cellular-level interactions, we established direct co-culture systems with isolated OA chondrocytes. After 7 days of direct contact, chondrocytes co-cultured with hESC-Mac exhibited more intense Alcian blue staining, suggesting enhanced proteoglycan synthesis, compared to those with PBMC-Mac or untreated controls (**Fig. 3c, d**). This suggested that direct cell-cell contact contributes to the restorative effects. We then asked whether soluble factors alone could mediate these benefits. Treatment of OA chondrocytes with conditioned medium from hESC-Mac cultures recapitulated the robust anabolic response, yielding significantly stronger Alcian blue staining than parallel treatments with PBMC-Mac conditioned medium (**Fig. 3e, f**), This confirmed that potent paracrine activity represents a key mechanism through which these macrophages promote cartilage repair. At the molecular level, we established a transwell co-culture system to analyze chondrocyte gene expression (**Fig. 3g**). hESC-Mac demonstrated remarkable capacity to reprogram the chondrocyte phenotype, significantly upregulating hyaline cartilage markers (COL2A1, ACAN) while suppressing fibrocartilage (COL1A1) and hypertrophic (COL10A1) markers (**Fig. 3h, i**). Notably, they also significantly reduced expression of cellular senescence markers p16INK4a and p21Cip1 (**Fig. 3j**), suggesting an additional role in mitigating chondrocyte aging.

**Figure 3.**
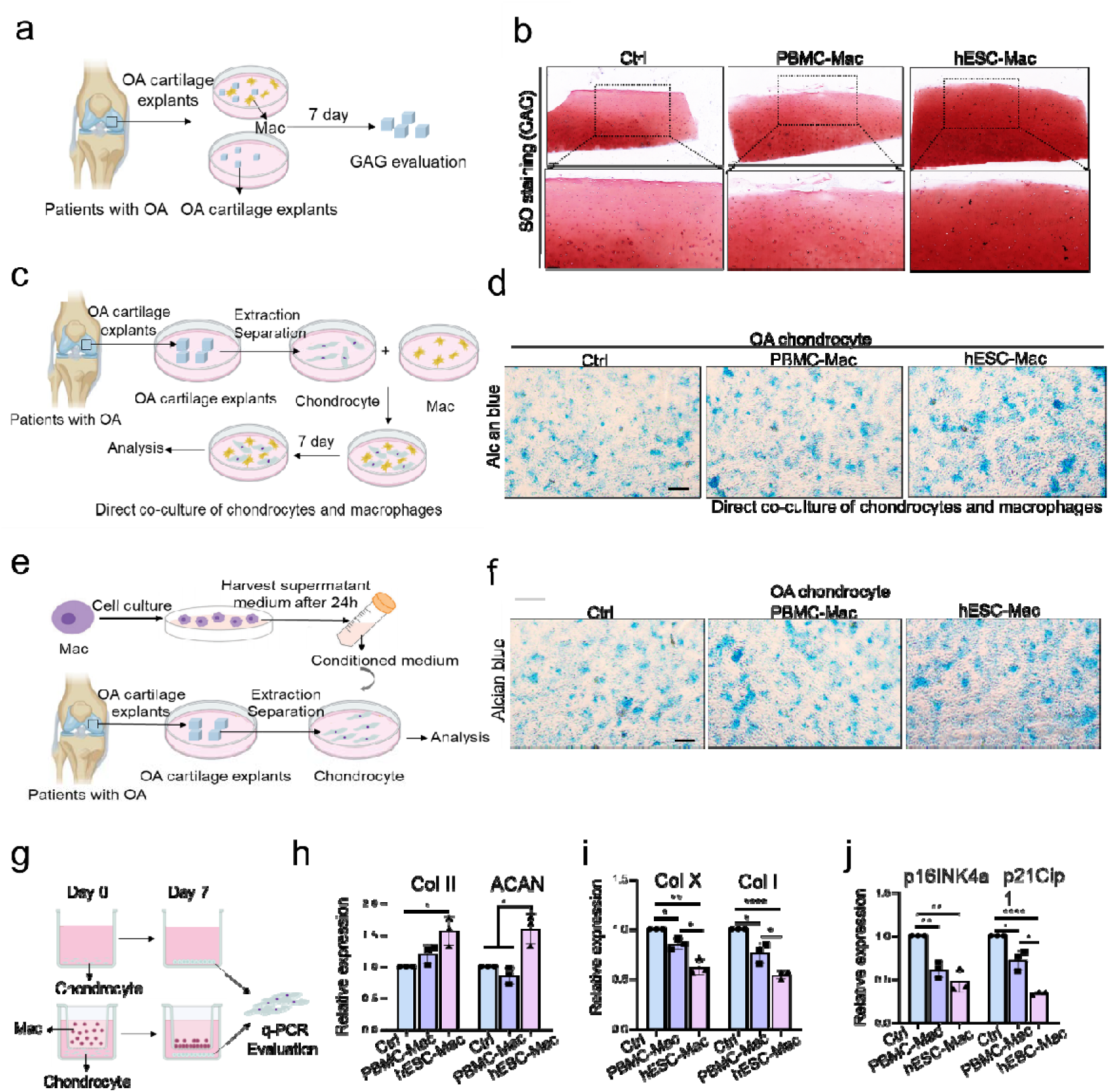
hESC-Mac demonstrates chondroprotective effects in human OA cartilage systems through direct contact and paracrine mechanisms. (a) Schematic illustration of the experimental setup for co-culturing human OA cartilage explants with macrophages for 7 days, followed by glycosaminoglycan (GAG) evaluation. (b) Representative images of GAG staining. Scale bars, 200 μm (up) and 100 μm (below). (c) Schematic diagram illustrating the collection of chondrocytes from human OA cartilage explants for direct co-culture with macrophages. (d) Representative Alcian blue staining showing proteoglycan content in OA chondrocytes after 7 days of direct co-culture with PBMC-Mac or hESC-Mac, compared to control. Scale bars, 100 μm (e) Schematic diagram illustrating the collection of chondrocytes from human OA cartilage explants for co-culture with macrophages-conditioned medium. (f) Alcian blue staining of OA chondrocytes exposed to conditioned medium (CM) from PBMC-Mac or hESC-Mac cultures compared with control medium. Scale bars, 100□μm. (g) Schematic diagram of the transwell co-culture system used for molecular analysis of chondrocyte gene expression. PCR assessment of (h) hyaline cartilage marker Col II and ACAN (i) fibrocartilage marker Col I and hypertrophic marker Col X (j) senescence markers (p16INK4a and p21Cip1) in OA chondrocytes following transwell co-culture with PBMC-Mac or hESC-Mac. Data represent mean ± SEM. *P<0.05, **P<0.01, ***P<0.001, ****P<0.0001.

The therapeutic relevance of this macrophage population was further supported by experiments in a murine model, where hESC-Mac effectively counteracted IL-1β-induced chondrocyte damage and proteoglycan loss (**Supplementary Fig. 8a, b**), mirroring the protective effects observed in human OA systems and confirming cross-species conservation of mechanism.

Collectively, these findings demonstrate that pro-regenerative hESC-Mac exert robust chondroprotective and pro-regenerative effects through both direct contact and paracrine mechanisms, effectively promoting a healthy chondrocyte phenotype while inhibiting degenerative and senescent pathways—highlighting their strong therapeutic potential for OA treatment.

### Intra-articular administration of hESC-Mac mitigates cartilage destruction and matrix loss *in vivo*

Given the marked chondroprotective effects of hESC-Mac observed *in vitro*, we proceeded to assess their therapeutic efficacy in a surgical murine model of osteoarthritis (OA). OA was induced by destabilization of the medial meniscus (DMM), followed by intra-articular injections of hESC-Mac or PBS control commencing at 4 weeks post-surgery, with endpoint analysis at 8 weeks (**Fig. 4a**). To track the joint retention of the administered cells, we performed *in vivo* imaging following a single injection of DIR-labeled hESC-Mac into osteoarthritic joints. Fluorescent signals were detected with progressive decline, remaining visible for up to five days—a retention profile comparable to that reported for human mesenchymal stem cells in similar contexts (**Supplementary Fig. 9a**).

**Figure 4.**
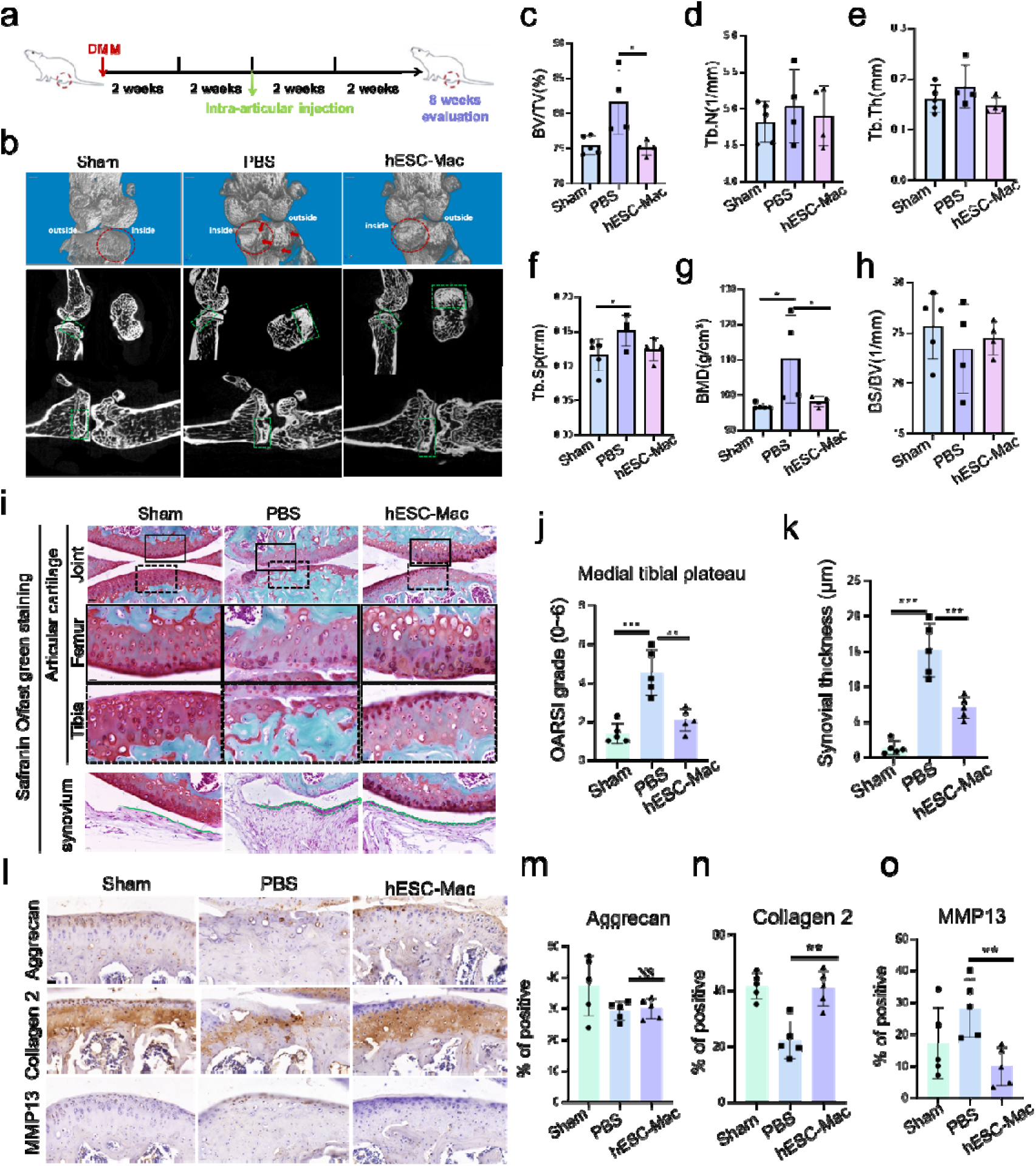
Intra-articular delivery of hESC-Mac mitigates OA progression and promotes cartilage repair *in vivo*. (a) Experimental timeline of the DMM-induced OA model and therapeutic intervention. Intra-articular injections of hESC-Mac or PBS control were administered at 4 weeks post-surgery, with endpoint analysis at 8 weeks. (n = 5 per group). (b) Representative micro-CT images of the medial tibial plateau from Sham, PBS-treated, and hESC-Mac-treated joints. Red circles indicate cartilage surface defects; Red arrowheads indicate severe damage to the cartilage; green boxes highlight the subchondral bone region of interest. Quantitative analysis of subchondral bone microarchitecture parameters: (c) BV/TV (Bone Volume/Total Volume, %), (d) Tb.N (Trabecular Number, mm□¹), (e) Tb.Th (Trabecular Thickness, mm), (f) Tb.Sp (Trabecular Separation, mm), (g) BMD (Bone Mineral Density, g/cm³), (h) BS/BV (Bone Surface/Bone Volume, 1/mm). (i) Representative images of safranin O/fast green staining of knee joint sections from each group. Boxed areas in the upper panels (overview of joint structure) are magnified below to show detailed cartilage structure (medial femur and tibial plateau) and synovial tissue. Scale bars: 50 μm (overview), 20 μm (magnified). (j) OARSI histological scores for cartilage degeneration, evaluated in medial femoral condyles and medial tibial plateaus. (k) Histological assessment of synovitis. Green lines outline the synovial lining thickness. (l) Representative immunohistochemical staining of aggrecan, collagen2, and MMP13 in articular cartilage across treatment groups. Scale bars, 20 μm. Quantitative analysis of the positive percentage of (m) aggrecan, (n) collagen2, and (o) MMP13 synovial thickness. Data are presented as mean ± SEM. *P<0.05, **P<0.01, ***P<0.001, ****P<0.0001.

Micro-computed tomography (micro-CT) analysis of the medial tibial plateau revealed substantial deterioration in PBS-treated OA joints, characterized by severe cartilage surface lesions (indicated by red arrows) and aberrant subchondral bone remodeling, evidenced by trabecular densification and loss of normal trabecular network architecture (**Fig. 4b**, green box). In stark contrast, joints treated with hESC-Mac displayed a marked preservation of cartilage integrity and subchondral bone microstructure, closely resembling sham-operated controls. Quantitative microarchitectural analysis demonstrated that hESC-Mac administration significantly normalized key bone parameters, notably reducing bone volume fraction (BV/TV, **Fig. 4c**) and bone mineral density (BMD, **Fig. 4g**) relative to the PBS-injected control. No statistically meaningful changes were found in trabecular number (Tb.N, **Fig. 4d**), trabecular thickness (Tb.Th, **Fig. 4e**), or bone surface-to-volume ratio (BS/BV, **Fig. 4h**) across the groups. While trabecular separation (Tb.Sp, **Fig. 4f**) was markedly elevated in the PBS group, hESC-Mac treatment not significantly prevented this aberrant increase.

Histological evaluation provided further evidence of cartilage protection. Safranin O/Fast Green staining confirmed the profound cartilage protection afforded by macrophage treatment (**Fig. 4i**). PBS-treated DMM mice exhibited extensive cartilage erosion, proteoglycan loss, and structural disruption on both femoral condyles and tibial plateaus. Conversely, joints receiving hESC-Mac injections maintained near-normal cartilage architecture with intense proteoglycan staining and smooth surfaces. Semi-quantitative OARSI scoring verified significantly reduced cartilage degeneration in the treatment group (**Fig. 4j**), underscoring the capacity of hESC-Mac to counteract cartilage catabolism.

Beyond cartilage protection, we observed significant modulation of synovial pathology. Administration of hESC-Mac attenuated synovitis, as indicated by reduced synovial hyperplasia (green line, **Fig. 4k**). To investigate the immunological mechanisms underlying this protective effect, we performed immunofluorescence analysis of macrophage infiltration and polarization in the synovium (**Supplementary Fig. 9b**). Quantitative assessment revealed significantly increased macrophage infiltration in PBS-treated OA joints compared to both sham controls and hESC-Mac-treated joints (**Supplementary Fig. 9c**). Crucially, polarization analysis demonstrated a pronounced pro-inflammatory milieu in PBS controls, characterized by significantly higher CD68^+^CD86^+^ M1 macrophage prevalence. In contrast, joints receiving hESC-Mac exhibited a marked shift toward an anti-inflammatory phenotype, with substantially increased CD68^+^CD206^+^ M2 macrophage dominance (**Supplementary Fig. 9d**). These findings indicate that intra-articular delivery of hESC-Mac not only reduces overall macrophage infiltration but also fundamentally reprograms the synovial immune microenvironment from a pro-inflammatory to a pro-reparative state, establishing a critical mechanism for their therapeutic efficacy in osteoarthritis.

Immunohistochemical analysis elucidated the mechanisms underlying cartilage preservation (**Fig. 4l**). Except for the ECM component aggrecan (**Fig. 4m**), which showed no significant difference, hESC-Mac-treated joints exhibited significantly higher expression of type II collagen (**Fig. 4n**), alongside reduced expression of the matrix-degrading enzyme MMP13 (**Fig. 4o**), indicating successful restoration of the anabolic-catabolic balance in chondrocytes. This pattern directly mirrors the effects observed in our earlier co-culture experiments, thereby strengthening the translational relevance of these *in vitro* findings.

These *in vivo* findings robustly demonstrate that a single defined immune cell product, pro-regenerative hESC-Mac, not as passive cellular bandages but as active instructors of joint homeostasis. Their ability to simultaneously maintain cartilage matrix, and resolve synovial inflammation underscores their potential as a multifaceted disease-modifying macrophage immunotherapy for osteoarthritis.

### hESC-Mac activate a TNFAIP3-mediated chondroprotective program

After confirming the strong cartilage-protective activity of hESC-Mac both *in vitro* and *in vivo*, we next aimed to define the key molecular pathways responsible for their therapeutic benefit. To comprehensively characterize the transcriptomic reprogramming induced by these specialized macrophages, we performed mRNA sequencing on human OA cartilage explants following co-culture with either hESC-Mac or under control conditions (**Fig. 5a**). Principal component analysis revealed striking segregation between the treatment groups, indicating fundamental transcriptomic alterations induced by hESC-Mac exposure (**Fig. 5b, c and Supplementary Fig. 10a**). We next integrated our differential gene expression data (hESC-Mac-treated vs. untreated OA cartilage) with published pseudobulk RNA-seq data comparing OA versus normal cartilage ^25^. This sophisticated analysis identified a core set of 30 genes whose pathological expression patterns were specifically reversed by hESC-Mac treatment—16 that were elevated in OA but normalized following therapy, and 14 that were deficient in OA but rescued upon intervention (**Fig. 5d and Supplementary Fig. 10b**). Notably, among the normalized genes were established antagonists of cartilage integrity: ITIH5, which blunts TGF-β signaling; COMP fragments that perpetuate TLR-mediated inflammation; and SAA2, a potent activator of NF-κB-driven catabolism. Conversely, the rescued genes encoded key promoters of tissue regeneration, including the matrix component FBLN1, the NF-κB checkpoint TNFAIP3, and the immunomodulator CD24 (**Fig. 5e**). This precise reversal of OA-associated gene dysregulation suggested that hESC-Mac actively restore cartilage homeostasis rather than merely suppressing inflammation.

**Figure 5.**
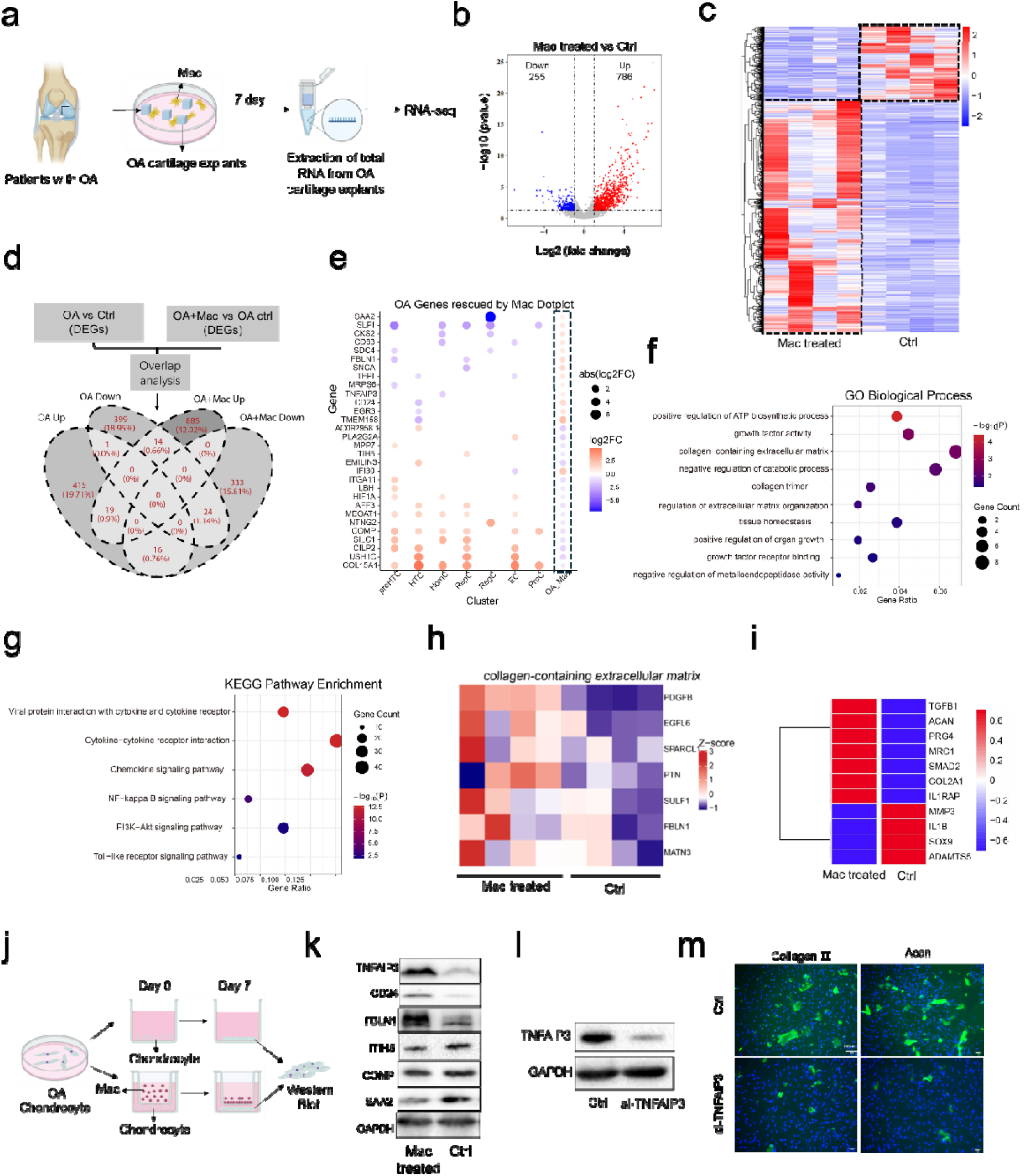
hESC-Mac reverse OA-associated transcriptomic alterations and activate a TNFAIP3-mediated chondroprotective program. (a) Experimental schematic of RNA sequencing analysis performed on human OA cartilage explants co-cultured with or without hESC-Mac for 7 days. (b) Volcano plot displays differentially expressed genes in hESC-Mac-treated versus control OA cartilage. Significantly upregulated (red) and downregulated (blue) genes are highlighted. (c) Heatmap displaying differentially expressed genes in hESC-Mac-treated versus control OA cartilage. (d) Venn diagram of OA-related differentially expressed genes (OA vs. normal cartilage in human articular cartilage cells) and genes rescued by hESC-Mac (OA+Mac vs. OA control). The intersection highlights DEGs whose expression is altered in OA and subsequently reversed following macrophage treatment. (e) Dotplot showing the rescued gene expression of OA-related DEGs and hESC-Mac–rescued DEGs. Articular cartilage cells group by subcell types, that is, preHTC (prehypertrophic chondrocyte), HTC (hypertrophic chondrocyte), HomC (homeostasis chondrocytes), RepC (reparative chondrocytes), RegC (regulator chondrocytes), EC (effector chondrocytes), ProC (proliferation chondrocytes). Only genes exhibiting opposite expression trends in the two comparisons (down in OA but up after macrophage treatment, or vice versa) are shown. Dot size and dot color reflect the log2FC values. These genes represent key pathological signatures whose dysregulation is functionally reversed by macrophage treatment. (f) Gene Ontology (GO) enrichment analysis of molecular functions, biological processes and Cellular Component modulated by hESC-Mac treatment in OA cartilage. (g) KEGG pathway enrichment analysis showing inflammatory signaling pathways suppressed by hESC-Mac treatment. (h) Heatmap visualization of collagen-containing extracellular matrix gene expression patterns in control versus hESC-Mac-treated OA cartilage. (i) Expression patterns of key chondrogenic markers and matrix-degrading enzymes in OA cartilage following hESC-Mac treatment. (j) Experimental workflow for protein validation using transwell co-culture of OA chondrocytes with hESC-Mac followed by western blot analysis. (k) Western blot analysis confirming reversal of pathological protein expression (ITIH5, COMP and SAA2) and enhancement of reparative factors (FBLN1, TNFAIP3 and CD24) in OA chondrocytes following pro-regenerative hESC-Mac co-culture. (l) Representative Western blot image showing TNFAIP3 protein expression in control (Ctrl) and TNFAIP3-knockdown (si-TNFAIP3) groups, with GAPDH as a loading control. (m) Representative immunofluorescence staining images of chondrocytes in Ctrl and si-TNFAIP3 groups for Collagen II and ACAN. Data are shown as mean ± SEM. *P<0.05, **P<0.01, ***P<0.001, ****P<0.0001.

Supporting this premise, gene ontology (GO) analysis revealed coordinated enrichment in processes fundamental to tissue repair, including collagen matrix assembly (collagen-containing extracellular matrix, collagen trimer, regulation of extracellular matrix organization and tissue homeostasis), growth factor responsiveness (growth factor activity and growth factor receptor binding), and restraint of metalloproteinase activity (negative regulation of metalloendo peptidase activity) (**Fig. 5f**). Concurrently, Kyoto Encyclopedia of Genes and Genomes (KEGG) pathway analysis indicated specific suppression of NF□κB and Toll-like receptor cascades (**Fig. 5g**), pathways centrally implicated in OA pathogenesis. We next focused on the collagen-containing extracellular matrix compartment, where hESC-Mac treatment robustly enhanced the expression of genes governing matrix synthesis and remodeling (**Fig. 5h**). This transcriptional reprogramming manifested functionally as coordinated upregulation of anabolic factors (TGFB1, ACAN, PRG4, COL2A1 and SOX9) alongside suppression of catabolic enzymes (ADAMTS5 and MMP3) (**Fig. 5i**), creating a molecular landscape favorable to cartilage regeneration.

To validate these findings at the protein level and establish direct mechanistic links, we employed transwell co-culture systems followed by western blot analysis (**Fig. 5j**). hESC-Mac consistently reduced pathological mediators (ITIH5, COMP and SAA2) while enhancing reparative factors (FBLN1, TNFAIP3 and CD24) in OA chondrocytes (**Fig. 5k and Supplementary Fig. 10c**). The central positioning of TNFAIP3 within this regulatory network prompted us to evaluate its functional necessity. we generated TNFAIP3-knockdown chondrocytes (**Fig. 5i and Supplementary Fig. 10d, e**). TNFAIP3 deficiency significantly attenuated Collagen - and aggrecan expression (**Fig. 5m**), confirming its essential role in mediating the chondroprotective effects of hESC-Mac.

Taken together, these multi-omics and functional studies demonstrate that hESC-Mac enact cartilage repair through a sophisticated regulatory network orchestrated around TNFAIP3. This molecular hub coordinates extracellular matrix restoration, constrains inflammatory signaling, and reestablishes chondrocyte homeostasis—providing a mechanistic foundation for their therapeutic efficacy in osteoarthritis.

## Discussion

Our findings establishe that embryonic skeletal development provides a master blueprint for regenerative macrophage therapy, revealing a targetable TNFAIP3-centered program that orchestrates chondroprotection in osteoarthritis. By integrating single-cell atlas of human embryonic skeletal development with functional validation^15^, we have uncovered a previously unappreciated heterogeneity of embryonic macrophages and their sequential emergence during chondrogenesis^26–28^. More importantly, this developmental insight directly inspired the design of a biomimetic induction platform that efficiently generates human embryonic stem cell-derived macrophages (hESC-Mac) with authentic regenerative properties.

A primary challenge in cell therapy is the precise definition of the desired therapeutic cell product. Our deconstruction of the embryonic macrophage landscape directly addressed this. We identified the LYVE1^+^ cluster emerges as a dominant and early population, strategically positioned to orchestrate skeletal morphogenesis through immunomodulatory and matrix-organizing functions **(Supplementary Fig. 1)**. Indeed, recent studies of LYVE1^+^ macrophages across diverse tissues reveal their conserved role as orchestrators of tissue development and repair^29^. Whether its mitigating inflammation in arthritic joints by producing specialized pro-resolving mediators, guiding stem cell fate decisions in adipose tissue through TGF-β signaling pathways^30^, or coordinating revascularization following cardiac injury through angiogenic factor secretion^31^. This subset, alongside other specialized populations like the chondrogenic COL9A1□ macrophages, provided a molecular benchmark for regeneration. Through systematic correlation analysis, we confirmed that our hESC-Mac most closely resembled LYVE1^+^ macrophages, sharing high expression of anti-inflammatory mediators (MRC1, CD163). Equally important was the shared transcriptional program with COL9A1□ macrophages, evidenced by co-expression of chondrogenic markers (ACAN, COL9A1, COL2A1, SOX9). This indicated that hESC-Mac possessed not only immunomodulatory capacity but also inherent pro-chondrogenic potential. Concurrently, we recognized HMGB1 as a defining feature spanning multiple embryonic macrophage subsets **(Fig. 2b)**, suggesting its role as a core regulator bridging immune function with regenerative programming. Traditionally viewed as a pro-inflammatory DAMP, HMGB1 exhibits remarkable functional pleiotropy in development, acting as a chemoattractant for stem cells and orchestrating tissue remodeling^32, 33^. The conserved high expression of HMGB1 in our hESC-Mac further validated their regenerative identity, signifying a programming for constructive tissue repair beyond mere immunosuppression^32^. Thus, we can defined our target therapeutic cell by the co-expression of MRC1—a marker of the predominant immunomodulatory LYVE1□ subset—and HMGB1, a pan-regenerative factor, dubbing them pro-regenerative macrophages.

A fundamental challenge in cell therapy is the reliable and scalable production of therapeutic cells. Our biomimetic dynamic platform directly addresses this challenge. By recapitulating key stages of primitive hematopoiesis and myelopoiesis, we achieve rapid, high-yield generation of hESC-Mac with exceptional purity and developmental fidelity. Mechanistically, we found that the enhanced efficiency of our system appears mediated through mechanosensitive pathways, as evidenced by upregulated β-catenin (CTNNB1)^34^, altered cytoskeletal dynamics (ACTB, RHOA and ROCK2), and enhanced nuclear translocation of the macrophage master regulator IRF8 **(Supplementary Fig. 4e)**. A detailed mechanistic exploration is beyond the scope of the current discussion. The successful establishment of this scalable production method is a crucial step towards the clinical translation of pro-regenerative macrophages-based immunotherapies, providing a reliable source of regenerative macrophages. Additionally, the low immunogenic profile of these cells, evidenced by minimal MHC class I and II expression^35, 36^. Studies have shown that certain embryonic and stem cell-derived populations exhibit intrinsic mechanisms for immune evasion, possibly reflecting a natural state prior to full immunological maturation^37–39^. While the precise mechanisms underlying this phenotype warrant further investigation, the resultant pro-regenerative macrophages exhibit a stable pro-regenerative phenotype, possess low immunogenicity, and demonstate functional competence, which make them ideal attributes for an allogeneic cell product **(Supplementary Fig. 4a)**. This scalable production method effectively solves the critical “cell source” problem, paving the way for clinical translation of regenerative macrophage therapies for OA other degenerative conditions.

Current OA therapies primarily aim to manage symptoms or delay structural deterioration, with limited regenerative capacity. While advanced approaches like MSC transplantation have demonstrated immunomodulatory potential, their capacity to drive robust and structural cartilage repair remains inconsistent, often failing to overcome the disease’s persistent inflammatory and catabolic milieu^40^. This limitation underscores the urgent need for novel therapeutic agents capable of actively reprogramming the joint microenvironment to achieve genuine regeneration. Our work introduces a paradigm shift by leveraging a defined macrophage as active therapeutic agent. We demonstrate that pro-regenerative macrophages are not mere anti-inflammatory vehicles but sophisticated instructors of joint repair **(Fig 2, 3 and 4)**. Their ability to simultaneously resolve synovial inflammation, promote anabolic matrix synthesis and reprogram the synovial immune microenvironment towards an anti-inflammatory state underscores their multifaceted disease-modifying potential. This positions macrophage immunotherapy as a promising new pillar for OA treatment, potentially complementing the existing strategies.

Mechanistically, transcriptomic and functional analyses revealed that pro-regenerative macrophages enact cartilage repair through a TNFAIP3-dependent program. This regulatory hub coordinates the reversal of OA-associated gene dysregulation, suppressing pathological mediators such as ITIH5, COMP fragments, and SAA2, while enhancing reparative factors including FBLN1 and CD24. The central role of TNFAIP3 was further underscored by loss-of-function experiments, in which its ablation in chondrocytes abrogated the induction of key matrix components, thereby dismantling a core mechanism of macrophage-mediated protection. This multi-level restoration of chondrocyte homeostasis—spanning extracellular matrix assembly, growth factor responsiveness, and restraint of NF-κB and TLR signaling—positions pro-regenerative macrophages as active instructors of joint repair rather than passive anti-inflammatory agents.

In summary, we propose a model wherein pro-regenerative macrophages deploy a TNFAIP3-centric regulatory network to recalibrate the OA joint environment. This network operates through a coordinated cascade: decommissioning pathological drivers (ITIH5, COMP and SAA2), enhancing structural resilience, and empowering TNFAIP3 to restrain inflammatory signaling, thereby re-establishing homeostasis. While our study identifies a potent therapeutic node and a scalable cell source, several aspects warrant further investigation. The precise mechanosensitive mechanisms driving efficient differentiation, the molecular basis of the low immunogenic profile, and the long-term fate of transplanted macrophages in more complex disease models are exciting avenues for future research. By bridging developmental biology with regenerative immunology, this work not only provides a transformative strategy for OA but also establishes a generalizable framework for developing macrophage-based therapies for a broad spectrum of degenerative diseases **(Fig 6)**.

**Figure 6.**
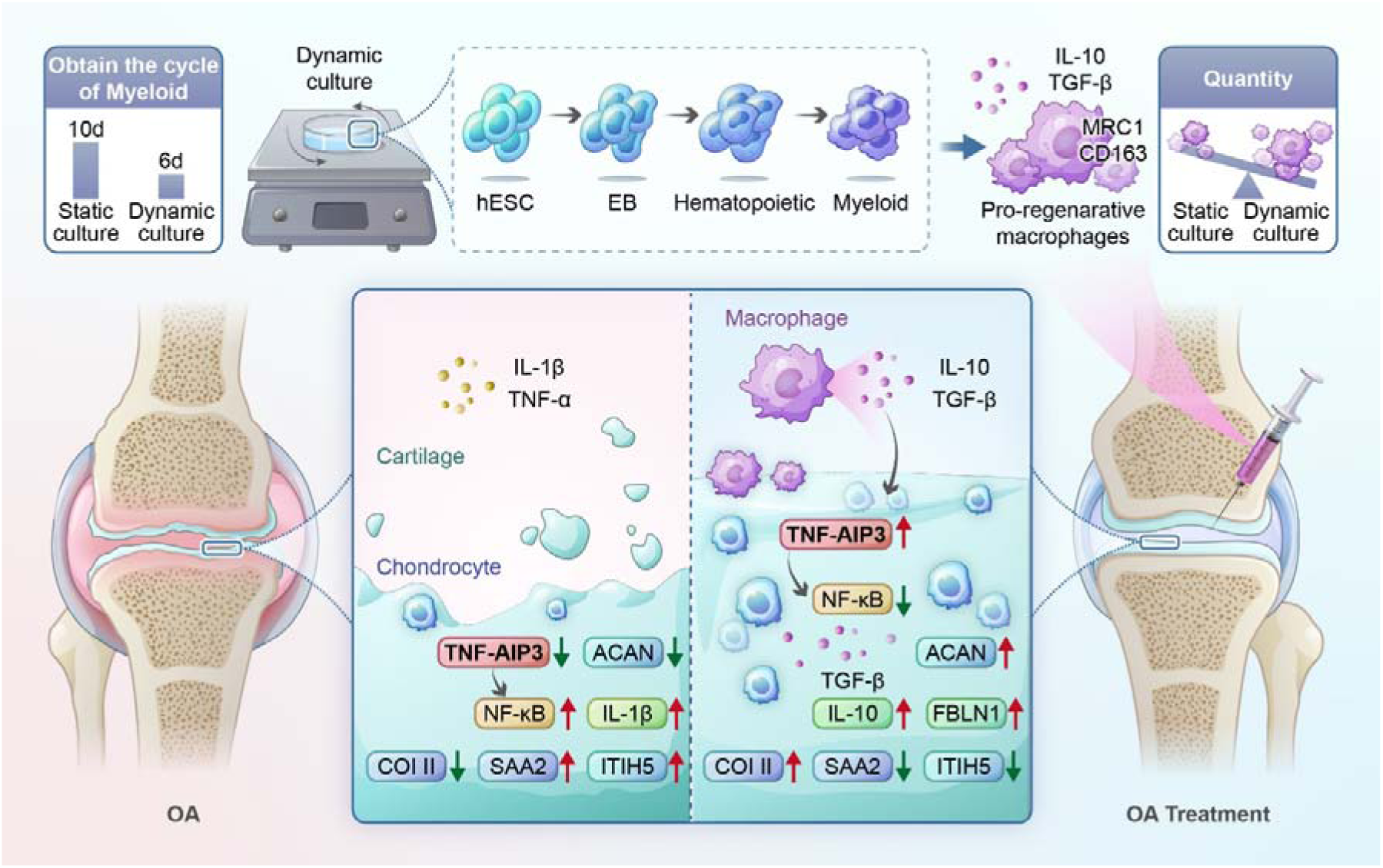
Schematic summary. Schematic of the optimized dynamic suspension culture system for generating pro-regenerative hESC-Mac from human embryonic stem cells (hESCs). Compared to conventional static culture, the dynamic (shaker) platform significantly shortens the differentiation timeline and enhances the yield and purity of the resulting macrophages. The therapeutic mechanism of pro-regenerative hESC-Mac in an osteoarthritic joint. These macrophages downregulate pathogenic factors (e.g., SAA2, COMP and ITIH5) and upregulate reparative mediators (e.g., FBLN1 and IL-10). Central to their chondroprotective effect is the activation of TNFAIP3 in chondrocytes. TNFAIP3 serves as a key molecular hub that suppresses NF-κB-driven inflammatory and catabolic signaling, promotes the expression of anabolic genes (e.g., COL2A1), thereby restoring cartilage homeostasis.

## Methods

### Ethical statement

All procedures involving human specimeg in Ehisiistudy were comauctedian compliance re reviewed and approved by the Ethics Committee of The Second Affiliated Hospital, Zhejiang University §qprool Of Medicine (Approval No. 2023LSYD0662). Human articular cartilage and synovial fluid were collected from the same institution, with donor information comprising three males and one female, averaging 65 years of age. The collection of samples and their subsequent experimental application were carried out in compliance with the ethical standards and authorized protocols established by the aforementioned committee.

### Macrophage differentiation from hESCs

Human ESCs were expanded on Matrigel-coated cultureware in mTeSR1 medium (85850, STEMCELL Technologies)^41^. To generate macrophages, we adapted a published differentiation protocol ^22^. Embryoid bodies (EBs) were formed and maintained under agitation from days 1 to 7. On day 1, EBs were treated with APEL II medium (STEMCELL Technologies) supplemented with BMP-4 and basic fibroblast growth factor (bFGF) to induce primitive streak-like mesoderm. From days 2 to 5, hematopoietic commitment was directed using APEL II medium with BMP-4, VEGF, bFGF and stem cell factor (SCF). Myeloid specification began on day 6 by adding bFGF, SCF, IGF-1, VEGF, GM-CSF, IL-3 and M-CSF to the medium. On day 7, Forty to fifty EBs were moved to a Matrigel-coated 6-well plate and cultivated in StemPro™-34 SFM medium (Gibco) with the identical cytokine cocktail (at the previously mentioned concentrations) up until day 17. Subsequently, from day 18, cells were matured for two days in a myeloid maturation medium with increased concentrations of GM-CSF and M-CSF. To promote terminal maturation, IL-3 was omitted from the myeloid maturation medium from day 19 onward. During the extended differentiation phase (days 7-24), non-adherent progenitor cells were collected at three-day intervals and replated onto fresh Matrigel-coated surfaces. The cells were then kept in X-VIVO medium (Lonza) that had 10% fetal bovine serum, M-CSF and GM-CSF.

### Chondrocyte isolation and culture

Human articular cartilage specimens were dissected into fragments of approximately 3-4 mm³. The fragments were subjected to sequential washes using Hank’s Balanced Salt Solution (HBSS) that included calcium and magnesium, followed by immersion in the same buffer. Tissue digestion was carried out by incubation with 0.5% (w/v) type II collagenase (Gibco, 17101-015) at 37°C overnight. To separate the liberated chondrocytes, the digested suspension was filtered using a sterile cell strainer.The remaining tissue remnants were subjected to a second instance of collagenase digestion, carried out under the same conditions. Chondrocytes were individually cultivated in a growth medium composed of DMEM/F12 (Gibco) enriched with 10% fetal bovine serum (Gibco) and 1% penicillin/streptomycin (Gibco). Based on the published methodology^42^.

### Generation of macrophages from human PB monocytes

Leucocyte concentrates were sourced from the Zhejiang Provincial Blood Center. PBMCs were extracted and then transformed into macrophages by cultivating them in RPMI-1640 medium supplemented with 50 ng/mL of human M-CSF and GM-CSF (PeproTech), following established protocols^43, 44^.

### Flow cytometry

Suspension cells, encompassing differentiated hESC-derived macrophages, were collected, rinsed with PBS, and then subjected to staining for 30 minutes at a temperature of 4°C using fluorochrome-conjugated antibodies that specifically target human CD14 (FITC).The expression of CD45 (FITC), HLA-ABC (APC), and HLA-DR/DP/DQ (APC) plays a crucial role in the immune response. After the staining process, cells underwent a washing procedure, were resuspended in PBS (phosphate-buffered saline), and subsequently analyzed using a BD flow cytometer. FlowJo software was employed to conduct data analysis. **Supplementary Table S2** contains a comprehensive list of antibodies.

### Quantitative real-time PCR (qRT-PCR) assay

RNA was extracted from specimens gathered at crucial phases throughout the hESC-to-macrophage differentiation process utilizing the RNeasy Mini Kit (Qiagen). DNase treatment using a DNA-free Kit (Qiagen) was performed to remove genomic DNA. The process of synthesizing the first strand of cDNA was performed using random hexamers and oligo (dT) primers, with the aid of Superscript III Reverse Transcriptase (Invitrogen). The synthesized cDNA was stored at a temperature of −20°C. Quantitative PCR reactions were prepared using FastStart Universal SYBR Green Master Mix (Roche) and executed on a QuantStudio™ 3 Real-Time PCR System (Thermo Fisher Scientific). To ensure consistency in cDNA input among reactions, the housekeeping gene β-actin was amplified simultaneously. The levels of gene expression were determined in comparison to β-actin through the application of the 2^−ΔΔCT^ method. In this research, a comprehensive list of all primer sequences employed can be found in **Supplementary Table S1**.

### Immunofluorescence staining

Following fixation, cells were subjected to permeabilization using 0.1% Triton X-100 for a duration of 5 minutes. To prevent non-specific binding, the sample was incubated for 20 minutes in a solution containing 5% bovine serum albumin (BSA). Mouse CD68, rabbit CD86, rabbit CD206, rabbit Collagen II, and rabbit Aggrecan primary antibodies were utilized and left to incubate overnight at a temperature of 4°C. On the subsequent day, cells underwent incubation using Alexa Fluor 488- or 594-conjugated goat anti-rabbit secondary antibodies that were diluted in PBS for a duration of 2 hours at room temperature. The nuclei underwent counterstaining using a concentration of 0.5 µg/mL DAPI for a duration of 5 minutes. A confocal laser scanning microscope was used to image stained samples. **Supplementary Table S2** contains detailed antibody information.

### Enzyme-linked immunosorbent assay (ELISA)

The macrophages were placed on plates at a concentration of 5×10□ cells per square centimeter and then allowed to grow for a duration of two days, after which the supernatant was collected and stored for analysis. Secreted levels of cytokines and chemokines, including tumor necrosis factor-α (TNF-α), interleukin-1β (IL-1β), IL-10, transforming growth factor-β (TGF-β), and CXCL18, were quantified using commercial ELISA kits purchased from MEIMIAN (China)

### Phagocytosis assay

To assess phagocytic activity, macrophages were incubated with 1.0 µm green fluorescent polystyrene microspheres (FluoSpheres™, Invitrogen, F13081) as per the manufacturer’s guidelines. After a 24-hour incubation, phagocytosis was evaluated by fluorescence microscopy or by quantifying bead uptake via live cell imaging.

### *In vitro* co-culture systems

Human cartilage tissues specimens were obtained with informed consent from patients undergoing total knee replacement surgery. Two co-culture platforms were established: For the OA cartilage explant-macrophage co-culture system, full-thickness cartilage discs (3-mm diameter) were maintained in X-VIVO medium (Lonza) as controls, while experimental groups involved direct contact co-culture with macrophages. The culture medium was maintained for 7 days with medium replenishment every 48 hours, followed by safranin O (SO) staining for histological evaluation. For the OA chondrocyte-macrophage co-culture system, three co-culture models were established: 1) Direct contact co-culture: chondrocytes and macrophages were seeded together in 12-well plates; 2) Transwell system: chondrocytes were seeded in the lower chamber while macrophages were placed in the upper chamber (0.4-μm pore size); 3) Conditioned medium experiments: Medium conditioned by macrophages for 24 hours was harvested and applied to chondrocyte monolayers. Co-cultures were maintained for 7 days with medium replenishment every 48 hours.

### Alcian blue staining

After the aspiration of the culture medium, the samples underwent three rinses with PBS and were then fixed using 4% paraformaldehyde for a duration of 30 minutes at room temperature. Following thorough PBS washes, samples were acidified by incubation in 0.1 N HCl for 5 minutes to achieve pH ∼1.0, ensuring specific binding to sulfated glycosaminoglycans. Subsequently, samples were immersed in a filtered 1% (w/v) Alcian Blue solution prepared in 3% acetic acid and stained overnight with gentle agitation. Non-specific background was removed through three 5-minute washes with 0.1 N HCl until clear visualization was achieved. All solutions were prepared using ultrapure water, with the staining solution filtered prior to use.

### Western blot analysis

The chondrocytes underwent a three-time rinse with ice-cold PBS and were then subjected to lysis on ice for a duration of 20 minutes utilizing RIPA buffer supplemented with a protease inhibitor cocktail. The process involves periodic vortexing, ensuring seamless connections between the words in short sentences while maintaining a similar meaning.To clear the cell lysates, they were subjected to centrifugation at 12,000 × g for a duration of 20 minutes while maintaining a temperature of 4°C. The liquid residue was gathered,The total protein concentration was determined using a BCA Protein Assay Kit (Thermo Fisher Scientific) and a bovine serum albumin (BSA) standard curve ranging from 0 to 2 µg/µL. The protein samples were then subjected to boiling at a temperature of 100°C for a duration of 5 minutes, utilizing the 5× Laemmli buffer. Equal protein quantities were resolved within 10% SDS-polyacrylamide gels at a constant current of 30 mA per gel. Separated proteins were then electrophoretically transferred onto nitrocellulose membranes using a wet transfer system under cold conditions. The membranes were impeded by immersing them in a solution comprising 5% non-fat dry milk, dissolved in Tris-buffered saline with Tween 20 (TBST), for a duration of two hours at ambient temperature. The sample was then left to incubate overnight at a temperature of 4°C, using specific primary antibodies.After TBST washes, the membranes were probed using suitable horseradish peroxidase (HRP)-conjugated secondary antibodies for a duration of one hour at room temperature. The Amersham Imager 680 RGB system (Cytiva) was employed to visualize immunoreactive bands. A list of all antibodies is available in **Supplementary Table S2**.

### Transfection

Chondrocytes were rinsed three times with ice-cold phosphate-buffered saline (PBS) and lysed on ice for 20 minutes using RIPA buffer containing a protease inhibitor cocktail, with periodic vortexing. Cell lysates were cleared by centrifugation at 12,000 × g for 20 minutes at 4°C. The supernatant was collected, and total protein concentration was determined with a BCA Protein Assay Kit (Thermo Fisher Scientific) using a bovine serum albumin (BSA) standard curve (0–2 µg/µL). Protein samples were subsequently denatured by boiling at 100°C for 5 minutes in 5× Laemmli buffer. Equal protein quantities were resolved with 10% SDS-polyacrylamide gels at a constant current of 30 mA per gel. Separated proteins were then electrophoretically transferred onto nitrocellulose membranes using a wet transfer system under cold conditions. Membranes were blocked with 5% non-fat dry milk dissolved in Tris-buffered saline with Tween 20 (TBST) for 2 hours at room temperature, then incubated overnight at 4°C with specific primary antibodies. Following TBST washes, membranes were probed with appropriate horseradish peroxidase (HRP)-conjugated secondary antibodies for 1 hour at room temperature. Immunoreactive bands were visualized using an Amersham Imager 680 RGB system (Cytiva). A list of all antibodies is available in **Supplementary Table S3**.

### Single-cell RNA sequencing (scRNA-seq)

The cell suspensions were adjusted to a final concentration of 1 × 10□ cells/mL in PBS. Single-cell partitioning, barcoding, and the construction of libraries were carried out in accordance with the manufacturer’s guidelines for the GEXSCOPE® Single-Cell RNA Library Kit (Singleron Biotechnologies) on a Singleron Matrix® automated platform. Each library was measured, adjusted to a concentration of 4 ng/μL, and then combined with others for sequencing. The pooled libraries underwent sequencing on an Illumina NovaSeq 6000 instrument employing a 150 bp paired-end configuration.

### scRNA-seq analysis

Raw sequencing data were processed with CeleScope software (v1.15.0, Singleron Biotechnologies) using standard settings to generate gene expression matrices. Gene expression matrices were constructed by aggregating data from cell barcodes, unique molecular identifiers (UMIs), and gene counts. Following the initial steps, bioinformatic analyses were carried out in the R programming environment (version 4.3.3), mainly relying on the Seurat toolkit (version 5.2.1). A preliminary quality control screening was conducted to eliminate cells of subpar quality, taking into account gene count thresholds (1,000-8,000 for hESC-derived macrophages; 200–8,000 for iPSC- and PBMC-derived macrophages)^22, 45^ and a mitochondrial gene percentage threshold (<15%). Putative doublets were detected and filtered out using the scDblFinder package (v1.16.0). Multiple datasets were consolidated using Seurat’s merge function. The process of normalization was carried out using the NormalizeData function, after which highly variable genes were identified through FindVariableFeatures and the data underwent scaling with ScaleData.Dimensionality reduction was accomplished via the application of principal component analysis (PCA), The top 30 principal components were employed to create two-dimensional embeddings through the use of Uniform Manifold Approximation and Projection (UMAP). To reduce batch effects, a data integration approach was implemented utilizing the canonical correlation analysis (CCA) technique within Seurat’s IntegrateLayers function. Cell types were annotated automatically with the SingleR package (v2.4.1) referencing the Human Primary Cell Atlas (celldex v1.12.0), and cluster predictions were subsequently refined by manual inspection. Macrophage identity was confirmed by assessing established marker gene expression. Polarization states (M1/M2) were quantified using a module scoring approach (AddModuleScore) based on defined gene signatures (**Supplementary Table S4**).

### Differential abundance testing

The ArrayExpress repository, specifically under access E-MTAB-14385 (www.ebi.ac.uk/arrayexpress), provided data on human embryonic skeletal development that was used for profiling. The analysis of differential abundance was conducted employing miloR (version 1.10.0) (https://github.com/MarioniLab/miloR) in R. Seurat objects were converted to SingleCellExperiment format, and samples were grouped by developmental stage (5PCW vs 11PCW). A k-nearest neighborhoods graph (k = 20, d = 30) was constructed and overlapping neighborhoods were defined and counted across samples (brc_code). Differential abundance between stages was tested using testNhoods with graph-overlap FDR correction, and neighborhoods were annotated by cell type using annotate hoods.

### Trajectory inference and pseudotime analysis

Developmental trajectories were inferred with Monocle3 (version 1.3.7). Macrophage subsets were isolated from the integrated immune cell dataset and subjected to standard preprocessing, comprising normalization, highly variable gene selection, and dimensionality reduction to derive a stable low-dimensional representation. This processed dataset was then transferred into Monocle3, including its UMAP coordinates and cluster information to ensure consistency. Monocle3 was used to infer how cells are connected in developmental space and to build a trajectory structure. The trajectory root was defined based on the differential abundance testing results and *in vivo* developmental (pcw) stages. Pseudotime values were visualized and mapped back to the Seurat object for downstream visualization.

### Analysis of intercellular signaling networks

Intercellular signaling between immune and chondrocyte populations was investigated with CellChat (v2.1.2). Immune cell types were merged into two broad groups (Myeloid and Lymphoid), while chondrocyte and macrophage subtypes were kept separate. The analysis was restricted to secreted signaling and extracellular matrix–receptor interactions using the CellChat human database. CellChat identified expressed ligand–receptor pairs, estimated the likelihood of communication between cell types, and filtered out interactions supported by fewer than 10 cells. The results were aggregated into pathway-level signaling strengths. Communication patterns and major sending/receiving populations were summarized using heatmaps, network diagrams, and centrality analysis. Outgoing and incoming signaling activities were summarized as heatmaps for representative pathways.

### mRNA-seq data analysis

Total RNA was extracted using Trizol Reagent (Thermo Fisher Scientific) and then underwent DNase I treatment to remove any remaining genomic DNA. Following purification, messenger RNA was subsequently purified using the KAPA Stranded mRNA-Seq Kit (Roche Diagnostics GmbH), followed by library construction for RNA-Seq with the same kit adapted for Illumina platforms. Sequencing was conducted by Genefund Biotech (Shanghai, China) on an Illumina system under paired-end 2×150 bp mode in accordance with the manufacturer’s protocols.

### Processing and alignment of sequencing data

Raw sequencing reads were subjected to quality filtering via Fastp (v0.23.2) in order to eliminate adapter contamination, discard reads with a length shorter than 36 base pairs, and trim low-quality bases (parameters: The parameters for the process include a tail window size of 4, a mean quality threshold of 20, and the detection of adapters specifically for paired-end data with a length of 36. The clean reads obtained were subjected to evaluation via FastQC with the default settings. The human reference genome GRCh38 was aligned using HISAT2 (v2.1.0) with option--rna-strandness RF--dta).

### Differential expression analysis

The abundance of gene expression was quantified using FPKM units through StringTie v1.3.4d with parameters set to -e and --rf. The edgeR package (version 3.24.2) was utilized in R for conducting differential expression testing. To assess statistical significance, false discovery rate (FDR)-adjusted *p*-values were calculated. Genes were considered differentially expressed when they met both criteria: adjusted *p*-value < 0.05 and absolute log□ fold change ≥ 1. The process of visualizing results was carried out utilizing the ggplot2 package in R.

### Functional enrichment analysis

The gene functional annotations were sourced from Ensembl release 96. In order to determine biologically significant pathways, In R, the ClusterProfiler package (version 3.4.4) was utilized to perform Gene Ontology (GO) and Kyoto Encyclopedia of Genes and Genomes (KEGG) enrichment analyses. Genefund Biotech (based in Shanghai, China) carried out the complete bioinformatic workflow for this section.

### Mouse osteoarthritis model and *in vivo* treatments

All experiments involving animals were conducted in compliance with protocols that had been reviewed and approved by the Institutional Animal Care and Use Committee.Researchers induced osteoarthritis (OA) in 8-week-old male C57BL/6J mice through a surgical procedure known as destabilization of the medial meniscus (DMM).Four weeks following the DMM surgery, animals were administered weekly intra-articular injections over a span of four weeks. Treatment groups were administered 6×10□ hESC-Mac (in 10 µL PBS), while control groups received an equal volume of PBS alone. For *in vivo* cell tracking, macrophages were pre-labeled with a near-infrared DIR fluorescent dye and monitored longitudinally using an IVIS® Spectrum imaging system.

### High-resolution micro-computed tomography

For structural assessment of subchondral bone, knee joints were imaged using a SkyScan 1276 µCT scanner with an isotropic voxel resolution of 8.8 µm. Three-dimensional reconstructions were generated with NRecon v2.1.1.3. Subchondral bone microarchitecture parameters were analyzed in the medial tibial subchondral bone region using CTAn v1.20.3.0 software.

### Immunohistochemical analyses

The joint samples underwent fixation, decalcification, and embedding in paraffin blocks. Serial sections with a thickness of 5 µm were either stained using Safranin O/Fast Green for general histology purposes or employed for immunohistochemistry. Previously described methods were adapted to form the immunohistochemical protocol. Before applying the stain, Antigen retrieval was carried out by subjecting the sections to incubation with 0.1% pepsin (Nacalai Tesque) in 0.5 M acetic acid (Sigma-Aldrich) at a temperature of 37 °C for a duration of 1 hour within a humidified chamber. Following the process of deparaffinization and rehydration, the endogenous peroxidase activity was effectively suppressed by subjecting it to a 0.3% solution of hydrogen peroxide (H□O□; Wako Pure Chemical) in PBS. To reduce non-specific binding, a solution with 2% bovine serum albumin (BSA) was employed. The following components were used: Wako Pure Chemical provided 0.1% Tween-20 (Sigma-Aldrich) and 0.01% Triton X-100 (Wako Pure Chemical). The sections were subsequently treated with primary antibodies and left to incubate overnight at a temperature of 4°C. The main antibodies utilized encompassed a monoclonal antibody specifically targeting type II collagen (M2139; Santa Cruz Biotechnology) and a monoclonal anti-type X collagen antibody (X53; Invitrogen, a subsidiary of Thermo Fisher Scientific, offers a wide range of high-quality products for life science research. After PBS washes, the sections were subjected to a 60-minute incubation period with horseradish peroxidase-conjugated mouse IgGκ light-chain binding protein (m-IgGκ BP-HRP; Santa Cruz Biotechnology is a company that specializes in the development and production of various biotechnological products. Following further washing, immunoreactivity was made visible by incubating with DAB substrate (Roche Applied Science) for a duration of 5 to 15 minutes. Ultimately, the nuclei were stained with hematoxylin as a counterstain.

### Quantification and statistical analysis

Statistical analyses were carried out utilizing GraphPad Prism version 10.4. Results derived from separate studies are showcased as mean□±□standard deviation (SD) or standard error of the mean (SEM), as specified.The analysis of differences between two groups was conducted through the use of unpaired two-tailed Student’s t-tests, with a focus on achieving a larger change while maintaining the original meaning and ensuring an increase in the number of words without any reduction. The analysis of comparisons among three or more groups was conducted using one-way analysis of variance (ANOVA). A P-value below 0.05 was considered statistically significant, with the following notation used to denote significance levels: *P < 0.05, **P < 0.01, ***P < 0.001, and ****P < 0.0001.

## Data availability

Single-cell RNA-sequencing data for human yolk sac and placenta were downloaded from the ArrayExpress repository (www.ebi.ac.uk/arrayexpress) under accession codes E-MTAB-11673 and E-MTAB-6701, respectively. Bulk RNA-seq and scRNA-seq data are available for peer review under GEO number GSE312225, and GSE312434 respectively. To review GEO accession GSE312225:

Go to https://www.ncbi.nlm.nih.gov/geo/query/acc.cgi?acc=GSE312225

Enter token kvkzueckjbephax into the box.

To review GEO accession GSE312434:

Go to https://www.ncbi.nlm.nih.gov/geo/query/acc.cgi?acc=GSE312434

Enter token mxmzoycitrelxiv into the box. Additional data supporting the findings of this study are available from the corresponding author upon reasonable request.

## Acknowledgements

This work was supported by the National Key Research and Development Program of China (2023YFB3813000), National Natural Science Foundation of China grants (NO. T2121004, 82394441, and 82301016), Postdoctoral Fellowship Program of CPSF (GZC20251573), Zhejiang Provincial Natural Science Foundation Project (MS26H250007), Zhejiang Province Postdoctoral Research Excellence Funding Project (ZJ2025116) and the China Scholarship Council (CSC). The authors thank Jinghua Fang (The Second Affiliated Hospital of Zhejiang University) for providing the human cartilage samples and synovia. We thank Xiaohui Zou (The first Affiliated Hospital of Zhejiang University) for micro-CT.

## Author contributions

Zhumei Zhuang performed experiments work and wrote the manuscript; Zhumei Zhuang and Hongwei Ouyang designed the project; Zhumei Zhuang and Zicong Liu performed scRNA-seq and RNA-seq analysis, supervised by Hendrik Marks; Zhumei Zhuang, Hang Su, Wei Sun and Qiuwen Zhu completed animal experiments, the histological analysis and micro-CT analysis; Hongwei Ouyang, Hua Liu and Xianzhu Zhang revised the manuscript; Jingyi Xu helped in extracting primary peripheral blood mononuclear cells; all authors commented on the manuscript.

## Supplemental Figures

**Supplementary Figure 1.**
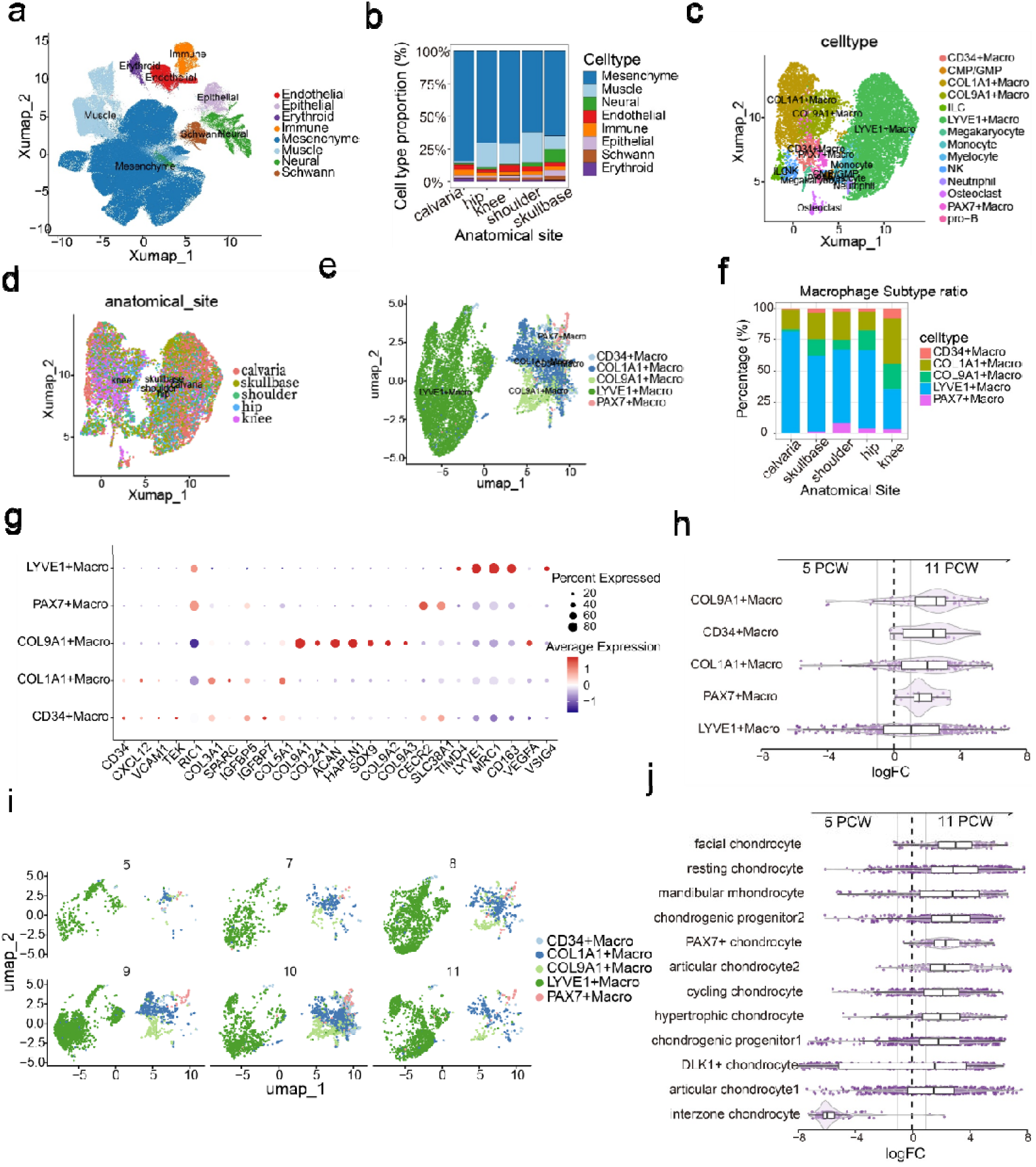
Single-cell profiling reveals the heterogeneity and developmental dynamics of embryonic skeletal macrophages. (a) Uniform Manifold Approximation and Projection (UMAP) visualization of the integrated single-cell transcriptomes (n=372,937 cells) from multiple anatomical sites (calvaria, skull base, shoulder, hip, knee), colored by major lineage annotations. (b) Relative cell-type abundance across anatomical locations. The bar plot of the proportion of the cell cluster compartment in each anatomical region sampled shows predominance of the mesenchyme compartment across anatomical regions. (c–d) UMAP systems of immune subpopulations, colored by cell type (c) and by anatomical origin (d). (e) Uniform manifold approximation and projection (UMAP) of macrophage subcell types from embryonic skeletal tissues across multiple developmental stages, colored by subset identity. (f) Distribution ratio of the five macrophage subsets across anatomical regions, demonstrating that LYVE1□ macrophages represent the dominant population at each skeletal site. (g) Dotplot showing the expression patterns of marker genes across five embryonic skeletal macrophage subsets. The color intensity represents the Z-score-scaled average expression of each gene, and the dot size indicates the percentage of cells expressing the gene. (h) MILO analysis of differential abundance of each macrophage subset across age (PCW), PCW<9 group to 5, PCW>9 group to 11. (i) UMAP depicting the dynamic changes of macrophage subsets across developmental stages (5 PCW to 11 PCW). (j) Developmental trajectory of chondrocyte lineages from 5 to 11 PCW, showing the progression from progenitor states (ChondroPro, Interzone chondrocyte) to specialized chondrocyte populations.

**Supplementary Figure 2.**
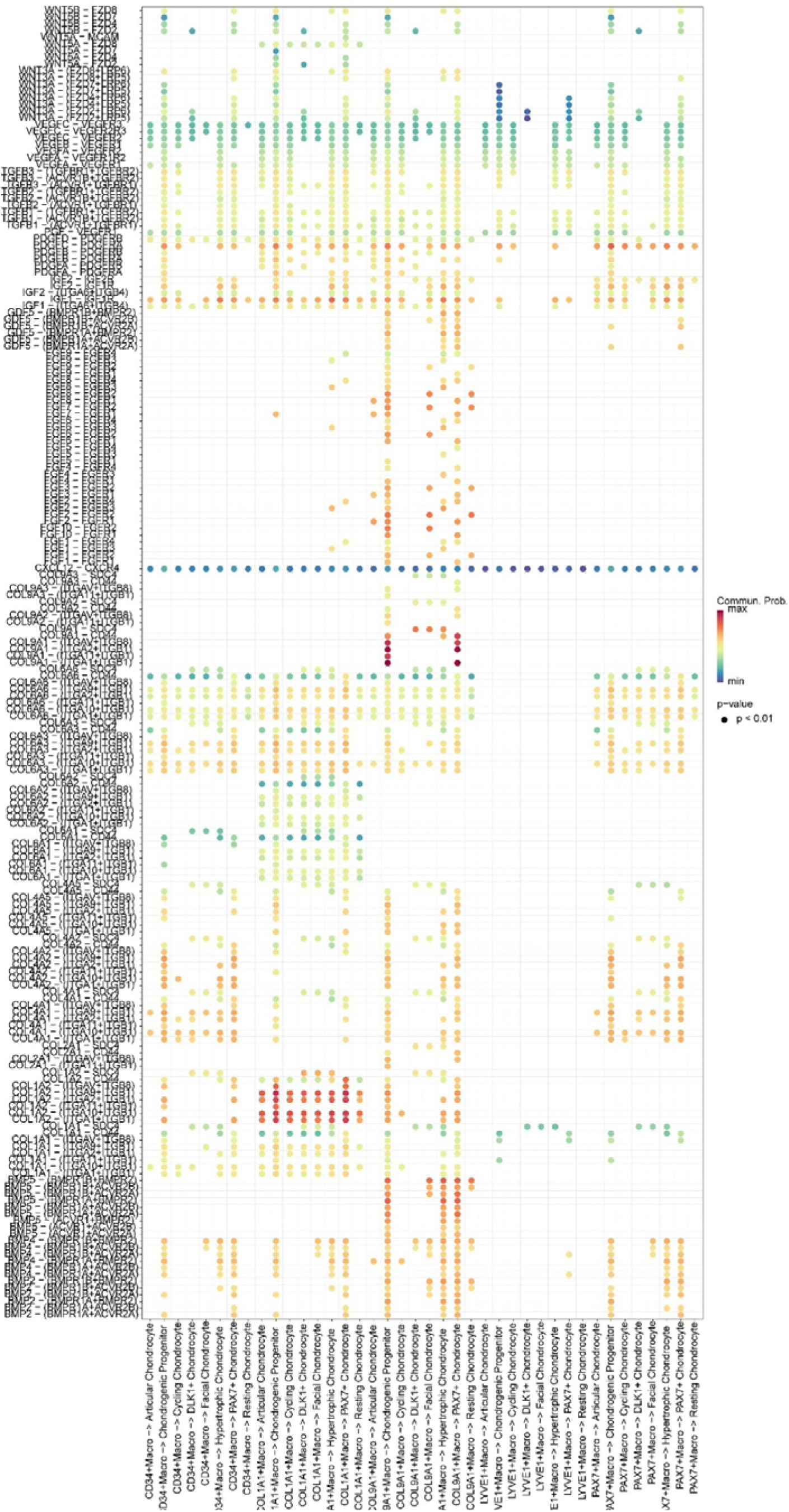
Embryonic skeletal macrophages establish comprehensive communication networks with chondrocyte lineages. Cell-cell communication analysis reveals extensive ligand-receptor interactions between specialized macrophage subsets (CD34^+^, COL1A1^+^, COL9A1^+^, LYVE1^+^ and PAX7^+^ macrophages) and multiple chondrocyte populations across differentiation states. The dot diagram illustrates specific signaling pathways from macrophage sources to chondrocyte targets, demonstrating that each macrophage subset engages in broad crosstalk with articular chondrocytes, chondrogenic progenitors, and specialized chondrocyte subtypes including cycling, DLK1^+^, facial, hypertrophic, PAX7^+^, and resting chondrocytes. This intricate network highlights the sophisticated regulatory capacity of embryonic macrophages in coordinating chondrogenesis and cartilage development.

**Supplementary Figure 3.**
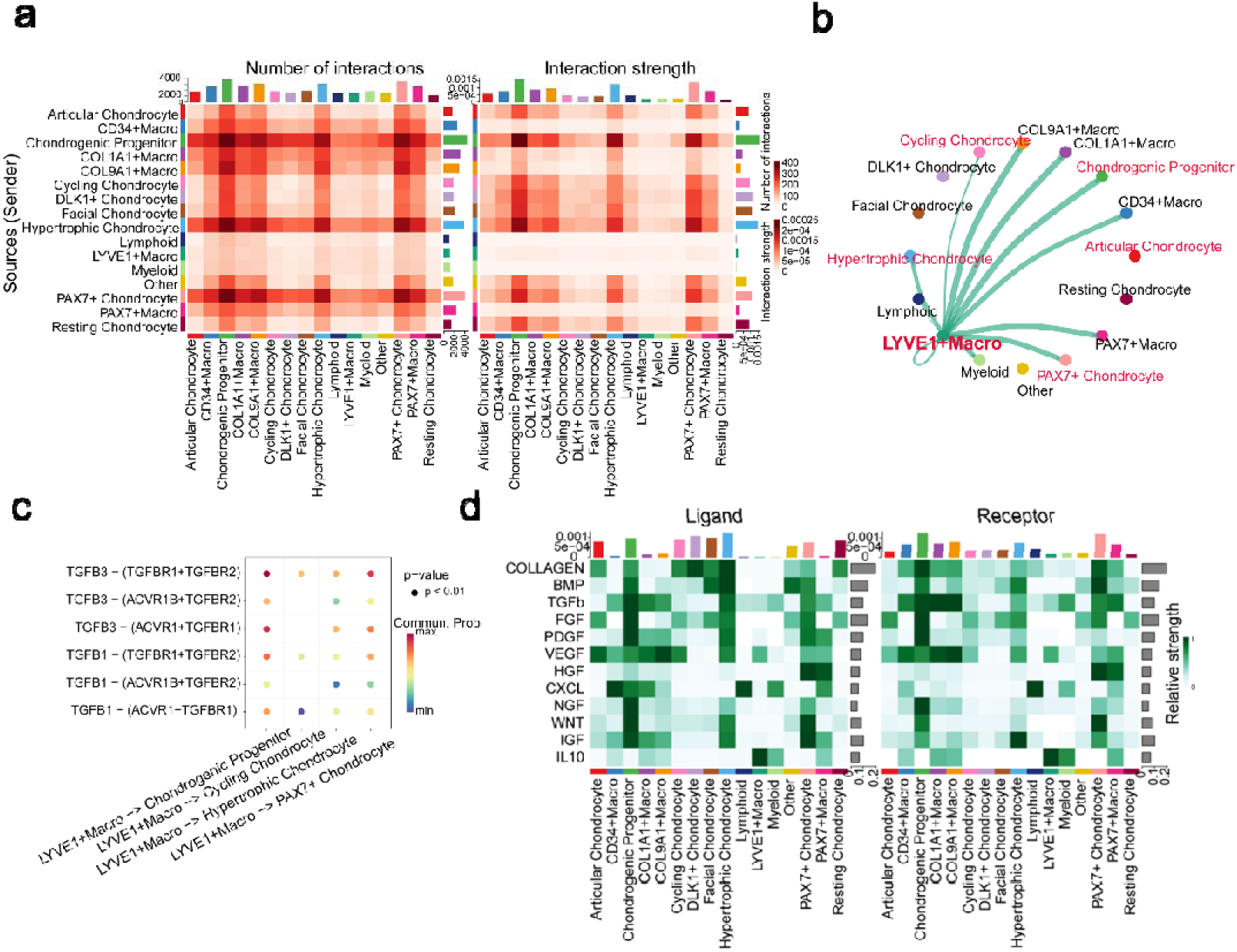
Cell-cell communication analysis reveals macrophage-chondrocyte interaction networks in embryonic skeletal development. (a) Heatmap visualization of interaction strength between source and target cell populations. Colors indicate communication probabilities, highlighting particularly robust interactions between specific macrophage subsets and cartilage-lineage cells. (b) Circos plot summarizing the LYVE1□ macrophages and other major cell types of communication network. Arcs represent cell populations, with connecting lines indicating significant interactions and line thickness corresponding to communication probability. LYVE1□ macrophages exhibit broad connectivity, with prominent interactions to chondrogenic progenitors and chondrocytes. (c) Selected TGF-β signaling pathways underlying LYVE1□ macrophage–chondrocyte interactions. Specific ligand–receptor pairs (e.g., TGFB1/B3 with TGFBR1/TGFBR2/ACVR1B) are significantly enriched (p < 0.01), indicating a dominant role for TGF-β in mediating macrophage–chondrocyte crosstalk during chondrogenesis. (d) Heatmap showing ligand (outgoing) and receptor (incoming) signals sent or received by immune and chondrocyte populations. Color intensity represents the relative interaction strength. Top barplots indicate the total outgoing (left) and incoming (right) signaling activity of each cell type, and side barplots show the overall strength of each signaling pathway across all cell populations.

**Supplementary Figure 4.**
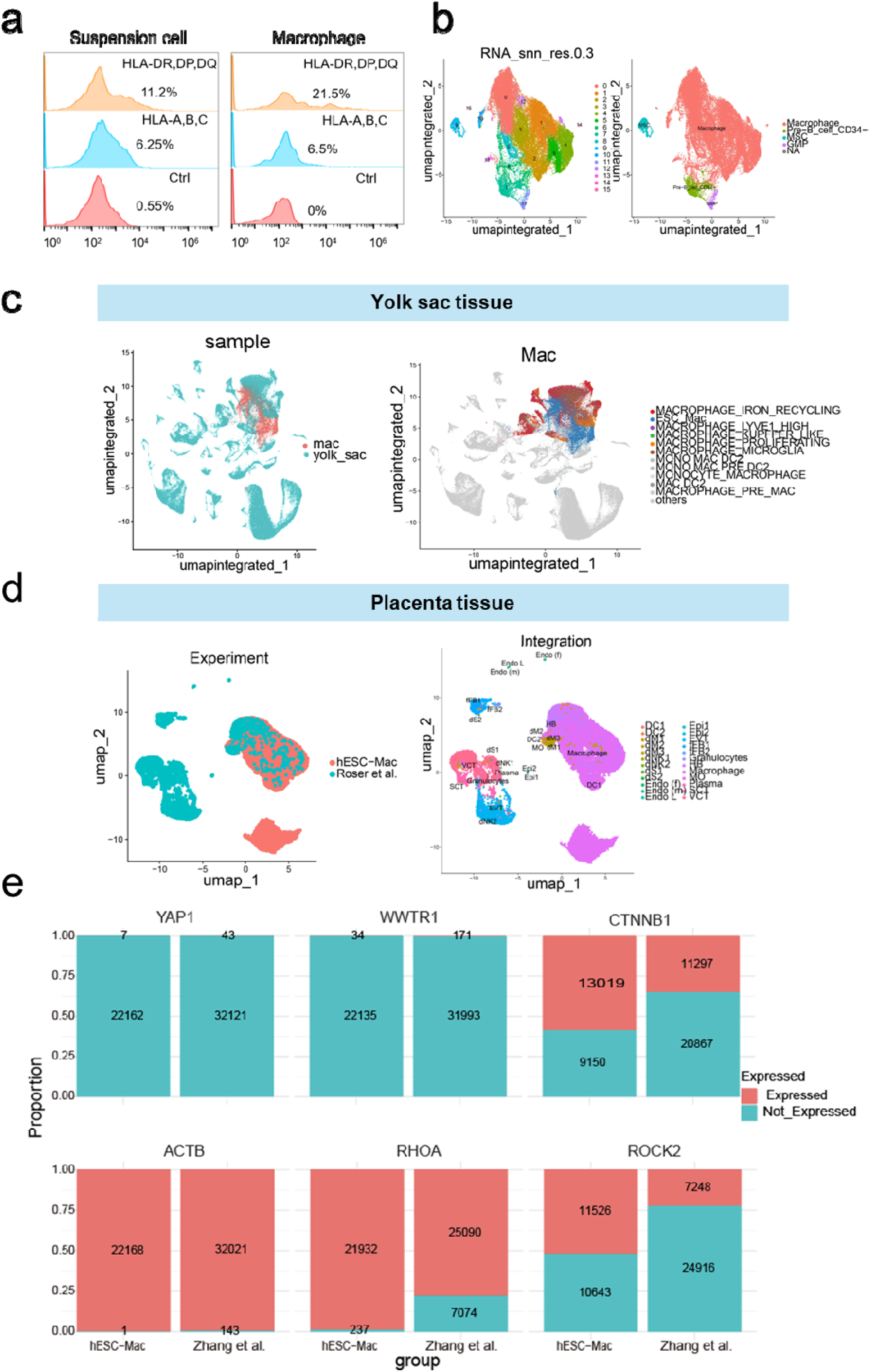
Immunogenicity and developmental origin of hESC-Mac. (a) Flow cytometry histograms showing the surface expression levels of antigen-presenting molecules HLA-DR, DP, DQ and HLA-A, B, C on undifferentiated hESC suspension cells and differentiated hESC-Mac. scRNA-seq analysis of hESC-Mac (b) UMAP plot of cells from the hESC-Mac culture, colored by unsupervised clusters (RNA_snn_res.0.3), revealing distinct cell populations. The analysis confirms the dominant presence of a Macrophage cluster, with minor populations of Pre-B_cell_CD34-cells and Granulocyte-Monocyte Progenitors (GMP). Integration of hESC-Mac scRNA-seq data with primary embryonic macrophage datasets to assess developmental origin. UMAP visualization of integrated single-cell transcriptomes from hESC-Mac and primary macrophages isolated from (c) yolk sac tissue and (d) placenta tissue samples. (e) Expression ratio of mechanosensitive gene expression under dynamic versus static induction conditions. Red color indicates expressed; blue color indicates not expressed. All data represent mean ± SEM.

**Supplementary Figure 5.**
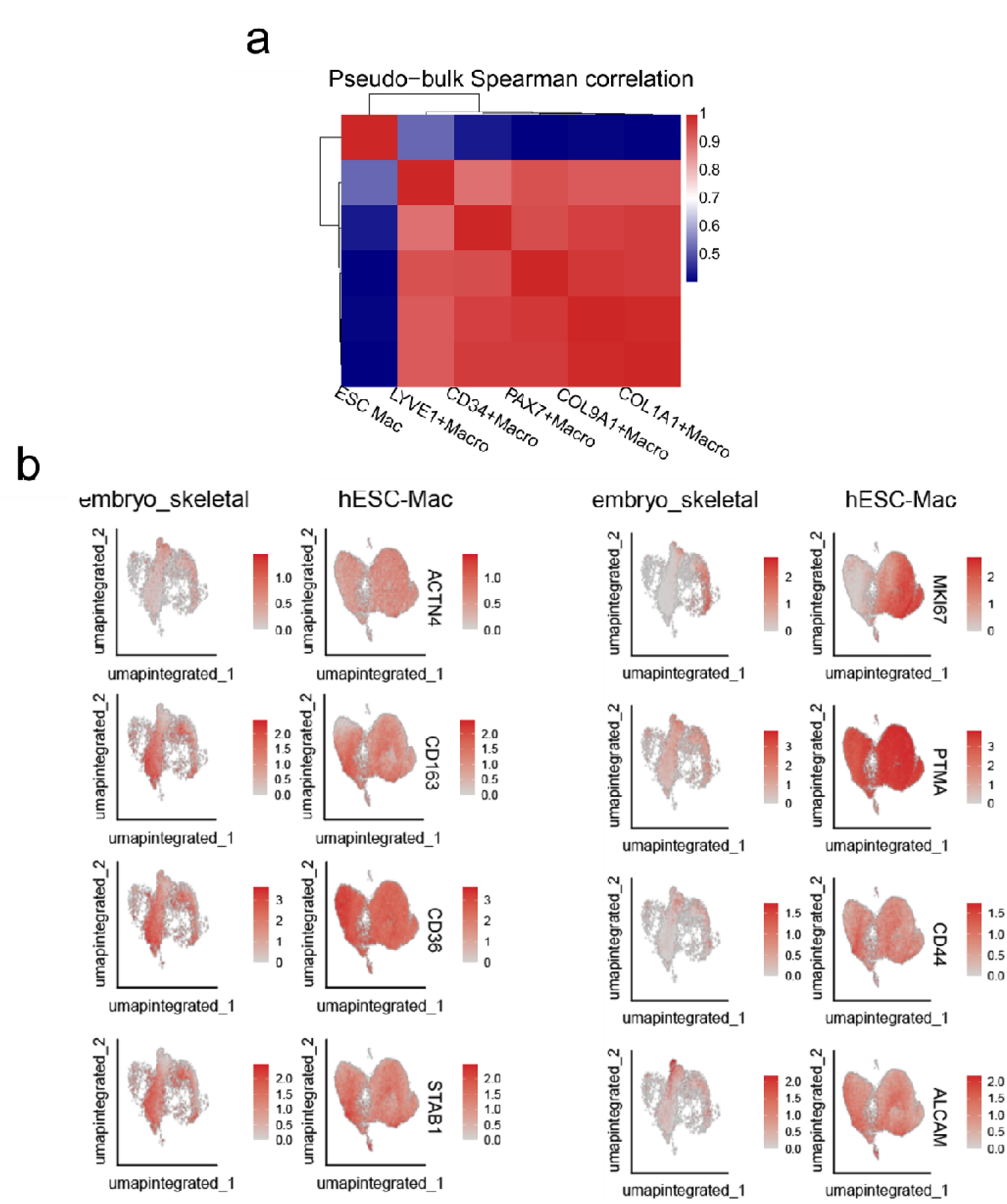
Integrated single-cell transcriptomic analysis reveals transcriptional similarity between hESC-Mac and embryonic skeletal macrophages. (a) Pseudo-bulk Spearman correlation analysis of hESC-Mac and macrophage subsets from embryonic skeletal tissue, correlation based on the expression of macrophage markers, markers from CellMarker 2.0. hESC-Mac shows the highest correlation with the LYVE1^+^ macrophage subset. Mac/Micro, Macrophage. (b) UMAP feature plots show the expression distribution of ACTN4, CD163, CD36, STAB1, MKI67, PTMA, CD44 and ALCAM in the integrated dataset of embryonic skeletal macrophages and hESC-Mac.

**Supplementary Figure 6.**
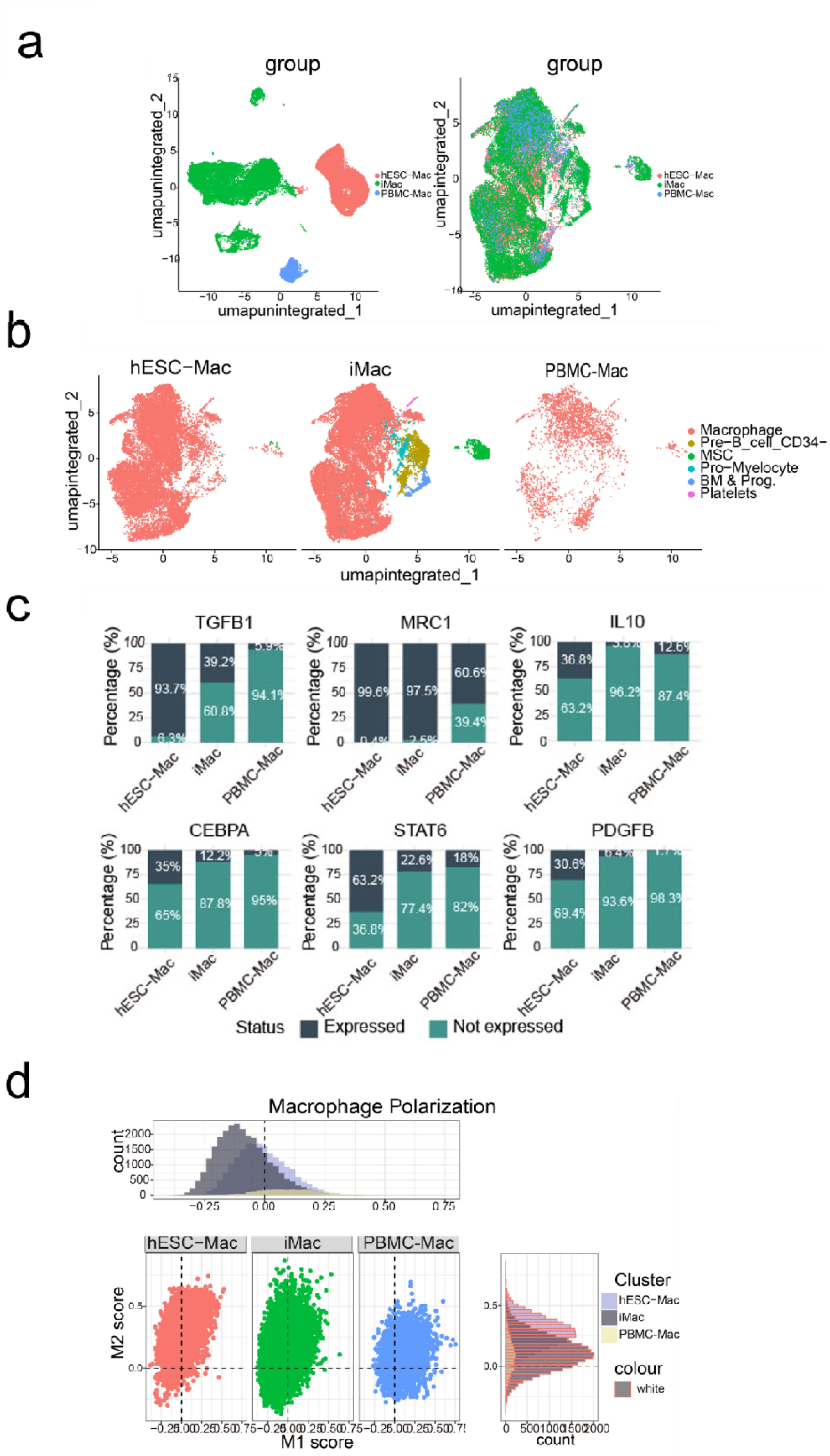
Comparative analysis of macrophage purity and gene expression across differentiation protocols. (a) Integration of single-cell RNA sequencing data from three macrophage sources: hESC-Mac (human embryonic stem cell-derived macrophages), iMac (macrophages derived using a static protocol), and PBMC-Mac (peripheral blood mononuclear cell-derived macrophages), colored by cell source. Left: unintegration UMAP, right: integration UMAP. (b) UMAP of cell annotation for (a), colored by cell type, revealing differences in population purity and the presence of non-target cell types (including Pre-B_cell_CD34-, MSC, Pro-Myelocyte, BM & Prog., and Platelets) across the different differentiation methods. (c) Bar graph showing the expression percentage of TGFB1, MRC1, IL10, CEBPA, STAT6, and PDGFB in macrophage cells of hESC-Mac, iMac, and PBMC-Mac populations as determined by scRNA-seq data. (d) Quantification of M1 and M2 polarization scores for hESC-Mac, iMac, and PBMC-Mac populations. hESC-Mac exhibit significantly elevated M2 scores coupled with the lowest M1 scores among all groups.

**Supplementary Figure 7.**
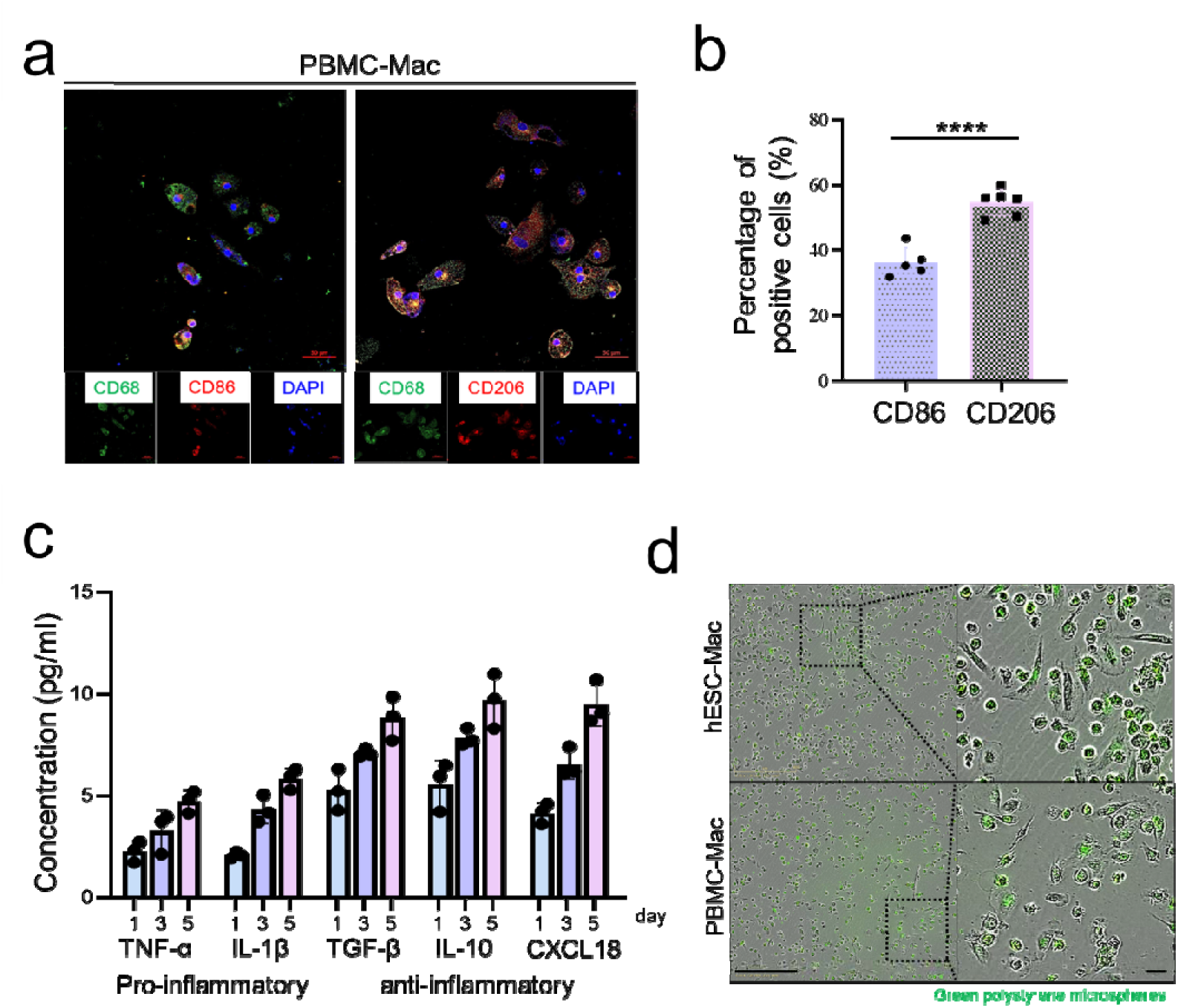
In vivo characterization the anti-inflammatory and phagocytic properties of PBMC-Mac. Representative immunofluorescence images (a) and corresponding positive quantification (b) of the anti-inflammatory marker CD206 (MRC1) and the pro-inflammatory marker CD86 in PBMC-Mac labeled with CD68. Scale bar, 50 μm. (C) quantitative analysis of the percentage of positive cells. (i) ELISA quantification of pro-inflammatory cytokines (TNF-α and IL-1β) and anti-inflammatory cytokine (TGF-β, IL10 and CXCL18) secretion in PBMC-Mac. (d) Phagocytosis assays demonstrate enhanced uptake of fluorescent beads by hESC-Mac. Scale bars: 400 μm (overview), 100 μm (magnified). All data represent mean ± SEM. *P<0.05; **P<0.01; ***P<0.001; ****P<0.0001.

**Supplementary Figure 8.**
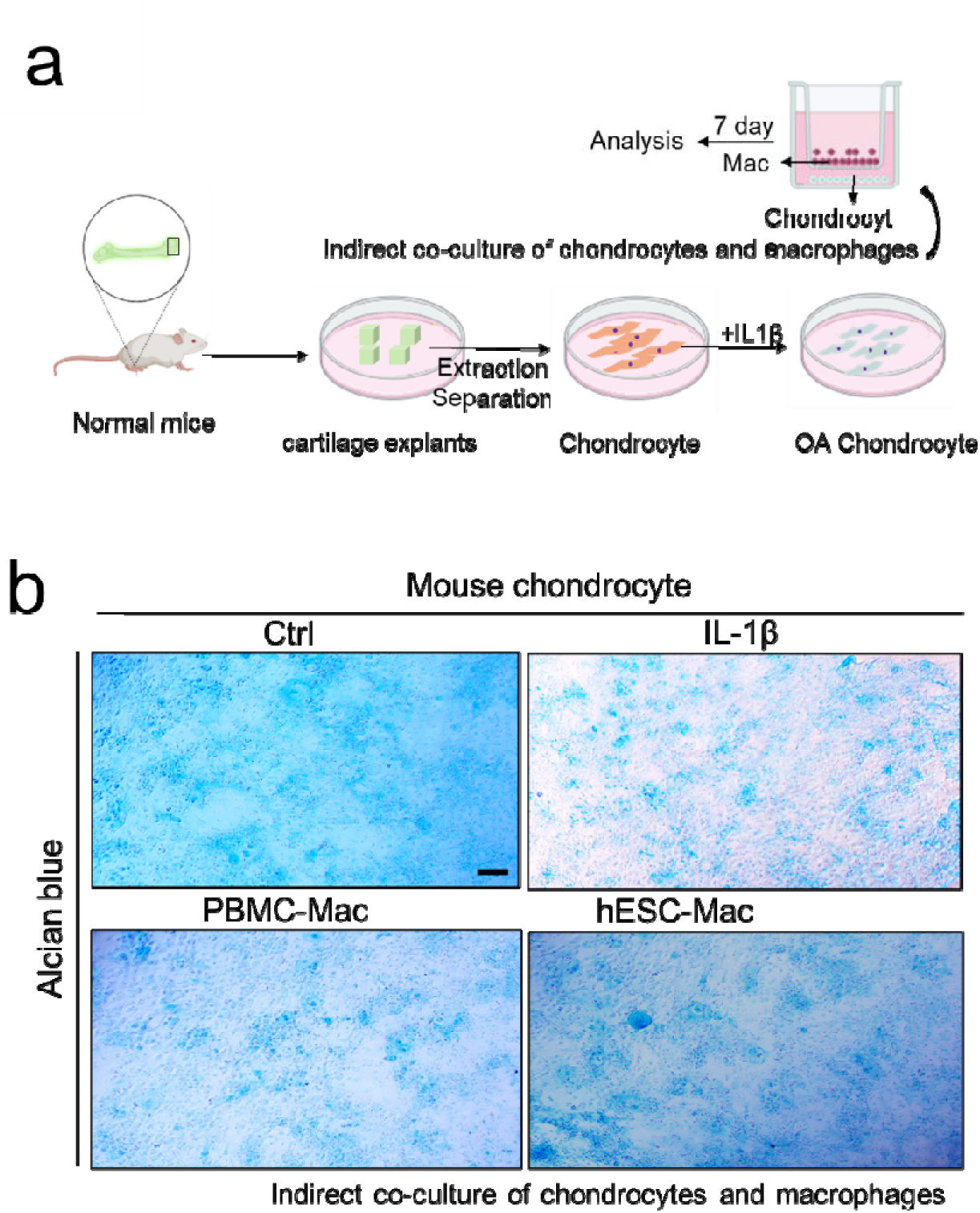
Protective effects of hESC-Mac in a murine IL-1β-induced chondrocyte damage model. (a) Schematic illustration of the experimental design using normal mouse cartilage explants. Chondrocytes were isolated and treated with IL-1β to create an in vitro OA model, and then co-cultured indirectly with macrophages using a Transwell system. (b) Representative images of Alcian blue staining showing proteoglycan content in murine chondrocytes under control conditions, after IL-1β treatment, and following intervention with macrophages.

**Supplementary Figure 9.**
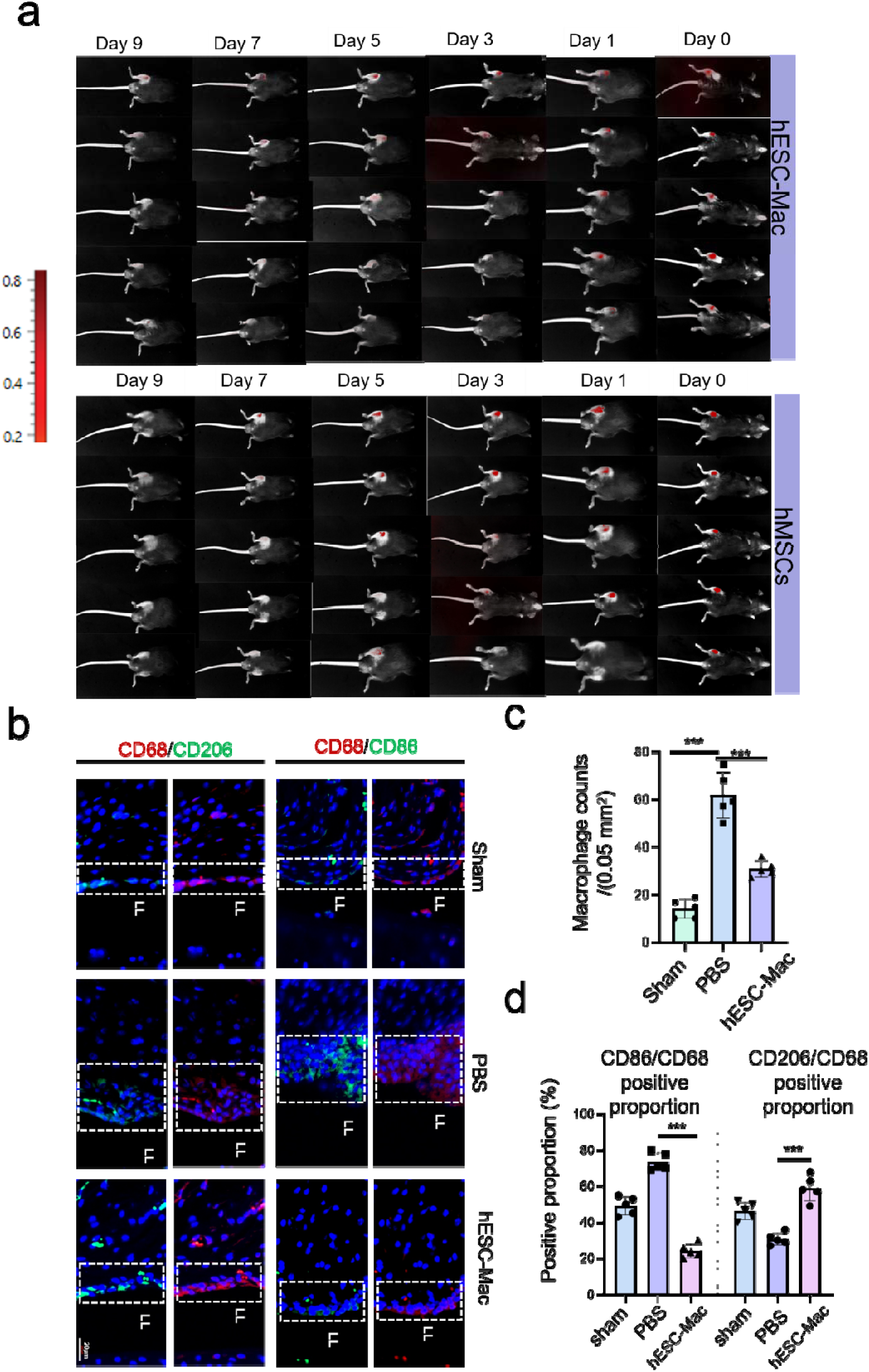
hESC-Mac reprogram the synovial immune microenvironment towards an anti-inflammatory state in OA joints. (a) In vivo fluorescence imaging of DIR-labeled hESC-Mac and hMSCs at various time points after intra-articular injection into OA joints. Signal intensity was monitored for up to 5 days. UMAP visualization of (b) Representative immunofluorescence images of synovial sections from Sham, PBS-treated, and hESC-Mac-treated DMM joints, stained for the macrophage marker CD68 (green), the M1 marker CD86 (red), the M2 marker CD206 (red), and DAPI (blue). Scale bar, 20 μm. (c) Quantification of total CD68^+^ macrophage infiltration in the synovium layer lining (LL). (d) Quantitative analysis of macrophage polarization states, presented as the percentage of CD68^+^CD86^+^ (M1) or CD68^+^CD206^+^ (M2) cells among total CD68^+^ macrophages. hESC-Mac treatment significantly shifts the balance towards an M2-dominant anti-inflammatory phenotype. All data represent mean ± SEM. *P<0.05; **P<0.01; ***P<0.001; ****P<0.0001.

**Supplementary Figure 10.**
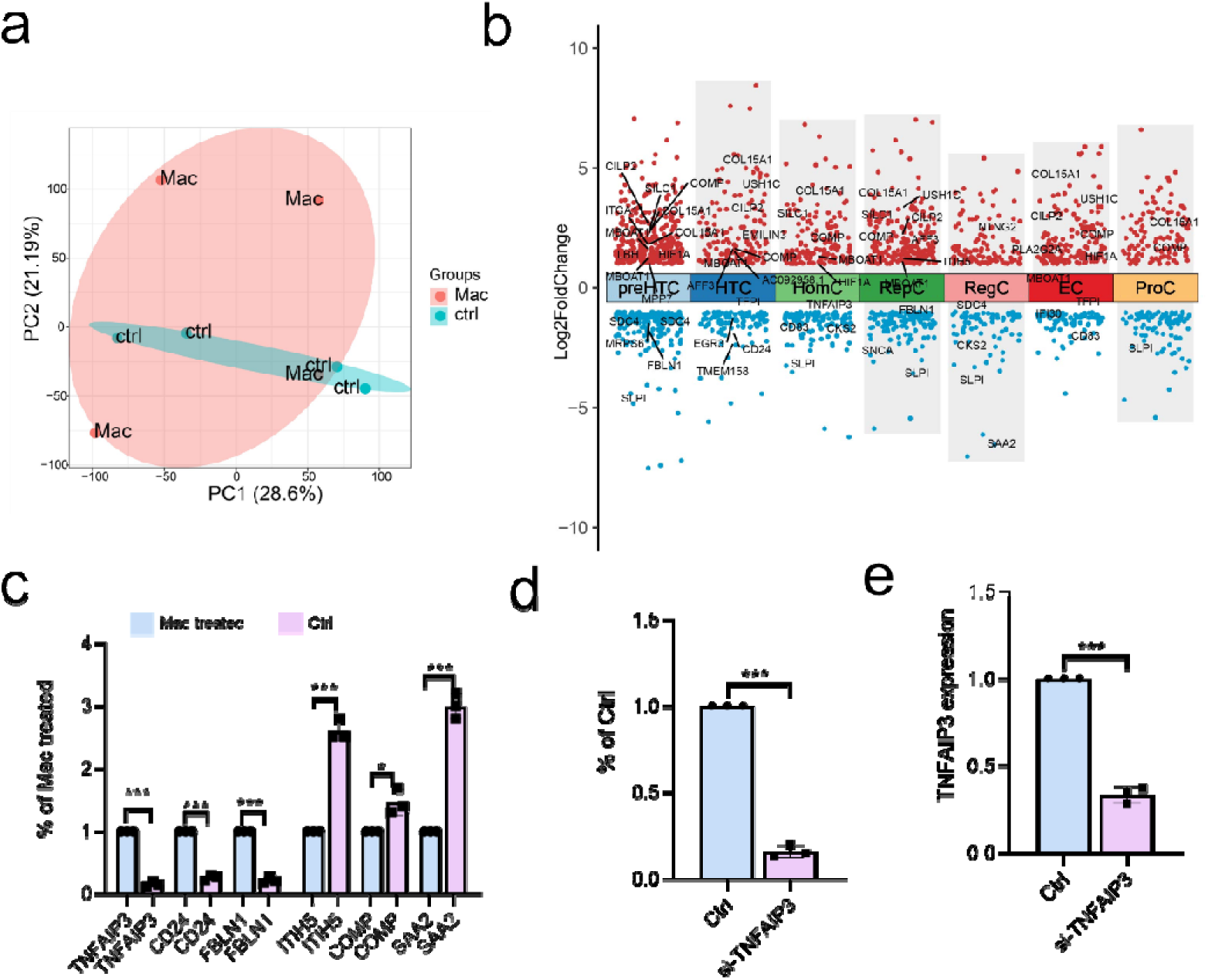
TNFAIP3 knockdown efficiency and its functional impact. (a) Principal component analysis (PCA) plot demonstrating distinct transcriptomic profiles between control and hESC-Mac-treated OA cartilage explants. (b) Dot plot of OA-related differentially expressed genes (OA vs. normal cartilage in human articular cartilage cell). Dot size represents absolute log2 fold change; color indicates direction of regulation. (c) Quantitative analysis of TNFAIP3, CD24, FBLN1, ITIH5, COMP, SAA2 protein levels from the Western blot in (Fig 6k). Data is normalized to GAPDH. (d) Quantitative analysis of TNFAIP3 protein levels from the Western blot in (d). Data is normalized to GAPDH. (e) mRNA expression levels of TNFAIP3 in Ctrl and si-TNFAIP3 groups. All data represent mean ± SEM. *P<0.05; **P<0.01; ***P<0.001; ****P<0.0001.

## Supplemental Tables

**Table S1.**
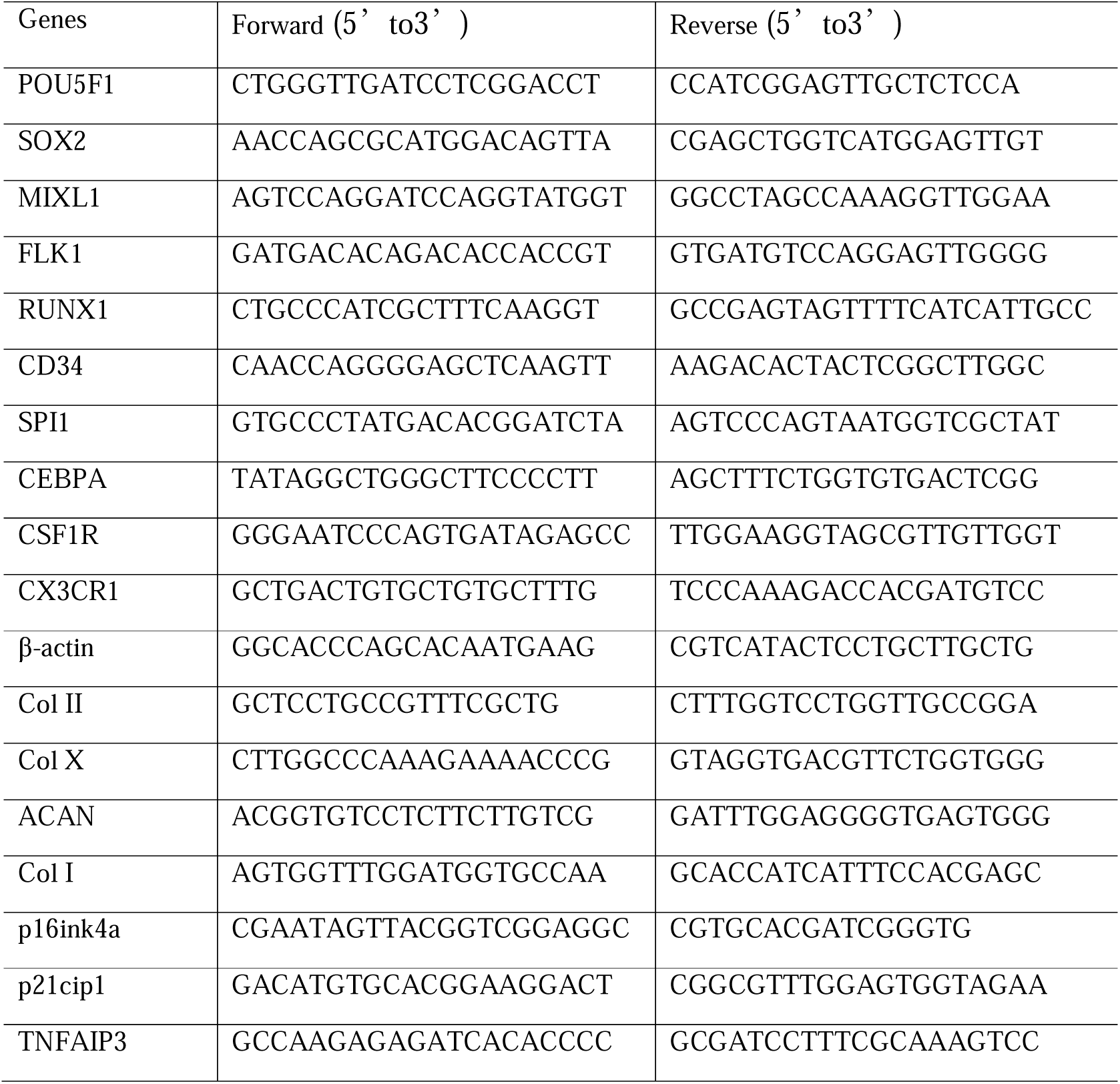
RT-PCR primers.

**Table S2.** Antibodies.

| Antibody | Company | Catalog | Use | Dilution Info |
| --- | --- | --- | --- | --- |
| FITC anti-human CD14 | BioLegend | 325603 | FC |  |
| FITC anti-human CD45 | BioLegend | 368507 | FC |  |
| FITC Mouse IgG1, $\kappa$ Isotype Ctrl | BioLegend | 400110 | FC | |
| APC anti-human HLA-A, B, C | BioLegend | 311409 | FC |  |
| APC anti-human HLA-DR, DP, DQ | BioLegend | 361713 | FC |  |
| APC Mouse IgG2a, $\kappa$ Isotype Ctrl | BioLegend | 400221 | FC | |
| Anti-rabbit CD86 | Proteintech | 13395-1-AP | IF | 1:500 |
| Anti-rabbit CD206 | Proteintech | 18704-1-AP | IF | 1:200 |
| Anti-mouse CD68 | Abcam | ab955 | IF | 1:50 |
| Goat Anti-Mouse IgG H&L (Alexa Fluor® 488) | Abcam | ab150113 | IF | 1:500 |
| Goat Anti-Rabbit IgG H&L (Alexa Fluor® 488) | Abcam | ab150077 | IF | 1:500 |
| Goat Anti-Rabbit IgG H&L (Alexa Fluor® 594) | Abcam | Ab150080 | IF | 1:500 |
| Anti-rabbit Acan | Abcam | ab315486 | IHC | 1:500 |
| Anti-rabbit Collagen 2 | Abcam | ab34712 | IHC | 1:200 |
| Anti-rabbit MMP13 | Abcam | ab39012 | IHC | 1:500 |
| Anti-rabbit Collagen 2 | Abcam | ab270993 | IF | 1:2000 |
| Anti-rabbit Acan | Servicebio | GB11373-100 | IF | 1:500 |
| Anti-mouse ITIH5 | Santa | sc-390885 | WB | 1:1000 |
| Anti-rabbit COMP | Abcam | ab74524 | WB | 1:1000 |
| Anti-rabbit SAA2 | Abcam | ab207445 | WB | 1:5000 |
| Anti-rabbit FBLN1 | Abcam | ab175204 | WB | 1:1000 |
| Anti-rabbit TNFAIP3 | Abcam | ab92324 | WB | 1:1000 |
| Anti-rabbit CD24 | Abcam | ab179821 | WB | 1:1000 |
| Goat Anti-Rabbit IgG H&L (HRP) | Abcam | ab6721 | WB | 1:2000 |
| Goat Anti-Mouse IgG H&L (HRP) | Abcam | ab6789 | WB | 1:2000 |

FC: Flow cytometry, IF: Immunofluorescence, WB: Western Blot

**Table S3.**
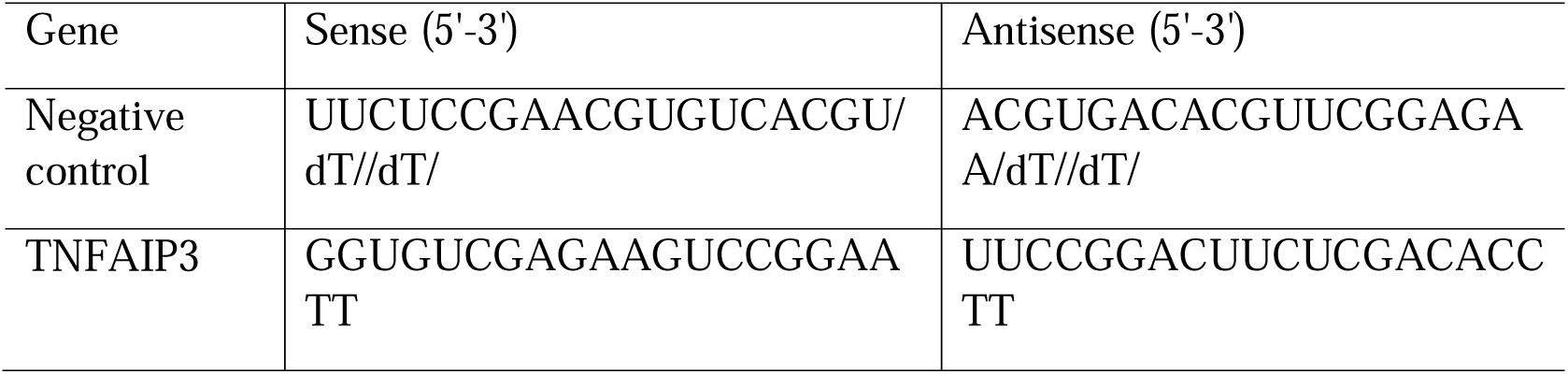
siRNA sequences.

**Table S4.**
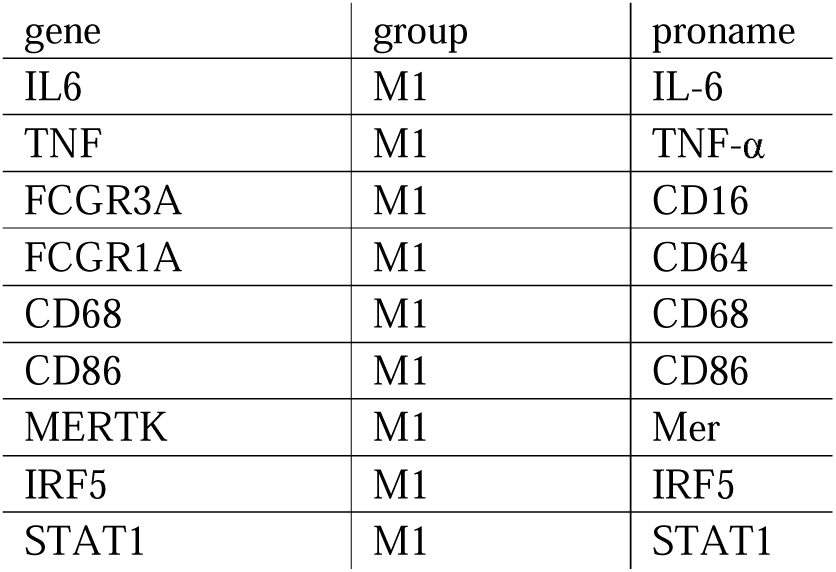
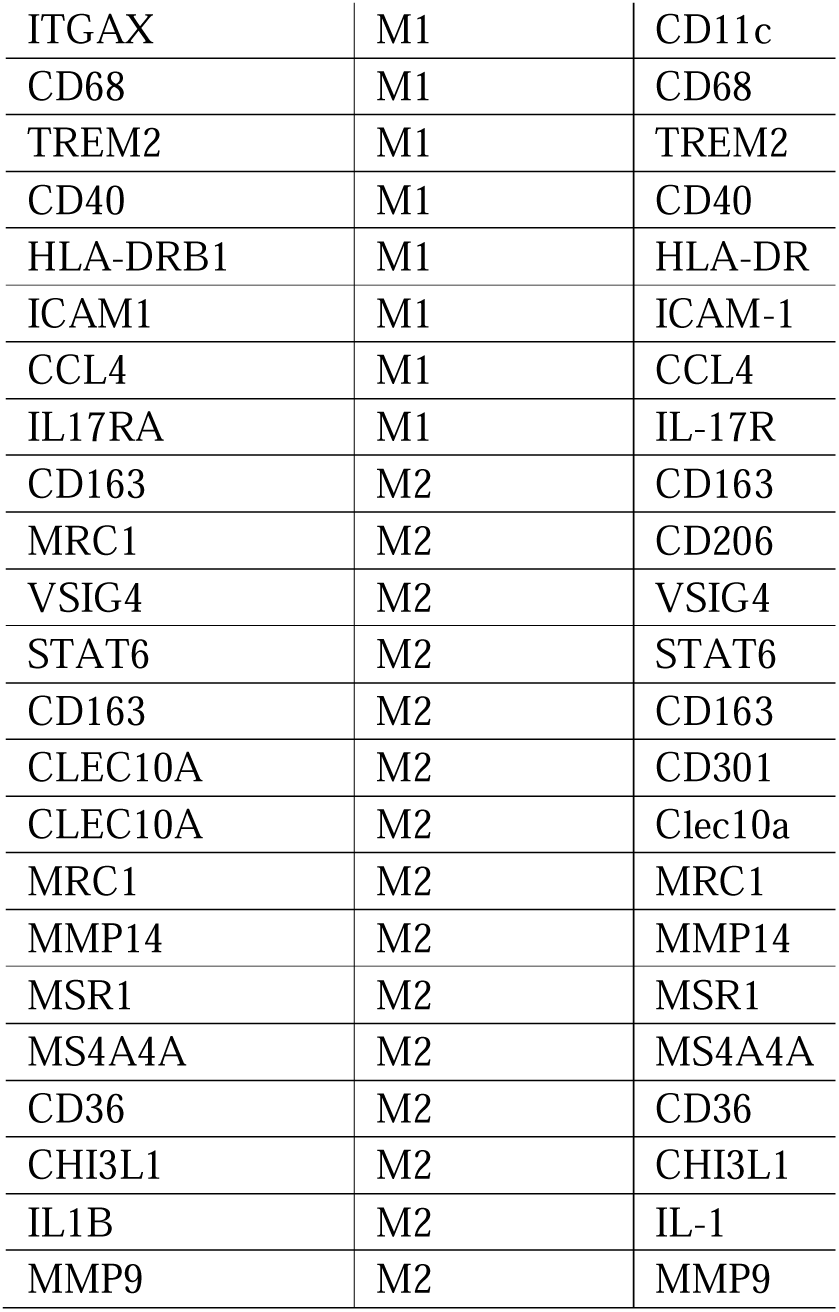
M1 and M2 markers.

